# Clathrin Light Chains are essential in negative regulation of cell death and immunity in Arabidopsis through interacting with autophagy pathway

**DOI:** 10.1101/2023.04.09.535952

**Authors:** Hu-Jiao Lan, Jie Ran, Lei Zhang, Ni-Ni Wu, Wen-Xu Wang, Min Ni, Ninghui Cheng, Paul A. Nakata, Jianwei Pan, Steven A. Whitham, Jian-Zhong Liu

## Abstract

Clathrin plays a critical role in clathrin-mediated endocytosis (CME) in plants, and it is required for autophagy in mammals. However, the functional interconnection of clathrin with autophagy has not been firmly established in plants. Here, we demonstrate that loss of function of clathrin light chain (CLC) subunit 2 and 3 results in salicylic acid (SA)- and H_2_O_2_-dependent accelerated senescence and activated defense responses in Arabidopsis, which are hallmarks of the autophagy-related gene (*ATG*) mutants. Similar to *atg* mutants, the *clc2-1clc3-1* double mutant has enhanced sensitivity to both carbon and nitrogen starvation and enhanced resistance to biotrophic bacterial and fungal pathogens. In addition, the autophagy flux was significantly reduced in the roots of *clc2-1clc3-1* mutant plants relative to Col-0 plants under carbon starvation conditions. Furthermore, our Yeast-2-hybrid (Y2H) and Luciferase complementation assays showed that CLC2 directly interacted with ATG8h and ATG8i. Mutations within the unique ATG8-interacting motif (AIM) of CLC2 as well as at the LIR/AIM-docking site (LDS) of ATG8h abolished the interaction between CLC2 and ATG8h. As anticipated, both GFP-ATG8h/GFP-ATG8i and CLC2 were subjected to autophagic degradation in the vacuoles. Together, our data revealed that the accelerated senescence and activated immune responses observed in Arabidopsis *clc2-1clc3-1* mutant plants result from impaired autophagy, and CLC2 participates in autophagy through direct interactions with ATG8h and ATG8i in an AIM1- and LDS-dependent manner. Our results unveil a previously unidentified link between the function of CLCs and autophagy.

## Introduction

Clathrin-mediated endocytosis (CME) plays critical roles in internalization and recycling of plasma membrane (PM)-localized proteins (Gelder et al., 2007; Sutter et al., 2007; Leborgne-Castel et al., 2008; Robert et al., 2010; Xu et al., 2010; Barberon, et al., 2011; Kitakura et al, 2011; Irani et al., 2012; Wang et al., 2013; Hao et al., 2014; Mbengue et al., 2016; Ortiz-Morea et al., 2016). Clathrin is a complex of triskelion shape consisting of three heavy chains and three light chains (Chen et al., 2011; McMahon et al., 2011). The *Arabidopsis thaliana* genome encodes two clathrin heavy chain (CHC) and three clathrin light chain (CLC) genes (Holstein, 2002; Chen et al., 2011). Loss of CLC2 and CLC3 leads to impaired auxin-regulated endocytosis of PIN proteins and consequently pleiotropic developmental defects including partial male sterility (Wang et al., 2013), altered blue light-triggered phototropic bending of the hypocotyl (Zhang et al., 2017) and impaired hypocotyl hook formation and opening (Yu et al., 2016). PM cargo proteins are usually mono-ubiquitinated prior to CME (Hicke and Dunn, 2003; Khaled et al., 2015). The mono-ubiquitinated proteins are firstly internalized in clathrin-coated vesicles (CCVs) to the trans-Golgi network (TGN)/early endosomes (EE), where endocytic and exocytotic/secretion pathways converge (Viotti et al., 2010). Subsequently, cargo proteins are sorted into the intralumenal vesicles (ILVs) of multivesicular bodies (MVBs) (Holstein, 2002; Tse et al., 2004; Viotti et al., 2010) and finally fused with vacuoles assisted by the endosomal sorting complex required for transport (ESCRT) machinery (Hwang and Robinson, 2009; Gao et al., 2014 and 2015; Khaled et al., 2015; Kolb et al., 2015; Zeng et al., 2019; Rodriguez-Furlan et al., 2019). Alternatively, these cargo proteins coated on the CCVs are deubiquitinated and recycled back to the PM from the TGN/EE (Khaled et al., 2015; Paez et al., 2016; Singh et al., 2018). Consistent with these observations, a recent study provided genetic evidence that clathrin function is not only required for endocytosis but also for exocytosis (Larson et al., 2017) and post-Golgi trafficking (Yan et al., 2021).

Macroautophagy (hereafter referred to as autophagy) is an evolutionarily conserved, dynamic catabolic process that engulfs damaged or no longer needed cytoplasmic components into double membrane vesicles called autophagosomes for vacuole/lysosome degradation (Li and Vierstra et al., 2012). Autophagic degradation can be either non-selective or selective. Under senescence and nutrient deprivation conditions, proteins, carbohydrates, and lipids can be broken down non-selectively via bulk autophagy to replenish the cells with carbon and nitrogen needed for survival and new growth (Li and Vierstra et al., 2012; Liu and Bassham, 2012; Boya et al., 2013; Teh and Hofius et al., 2014). On the other hand, aggregated proteins, damaged organelles induced under stress conditions, or even invading pathogens can be cleared selectively by autophagy, in which lipidated ATG8 proteins anchored on the autophagosome membrane recruit cargoes to autophagosomes through interacting with ATG8-interacting motif (AIM)-containing cargo receptors (the core AIM sequence is defined as W/F/Y-XX-L/I/V) (Li and Vierstra et al., 2012; Liu and Bassham, 2012; Noda et al., 2010; Ran et al., 2020). A LIR/AIM docking site (LDS) within the ATG8 amino acid sequence is responsible for interacting with cargo receptors (Marshall et al., 2019).

Nine ATG8 isoforms have been identified in Arabidopsis (Hanaoka et al., 2002; Kellner et al., 2017). Plant ATG8 genes are grouped into two clades by phylogenetic analysis (Kellner et al., 2017). The C-terminal Arg of newly synthesized ATG8 in clade I (ATG8a to ATG8g) is cleaved by ATG4, resulting in an exposed Gly residue at its C terminus. The lipid molecule phosphatidylethanolamine (PE) is then conjugated to the exposed Gly residue of ATG8 sequentially assisted by the E1-like enzyme, ATG7 and the E2-like enzyme ATG3, respectively (Ohsumi, 2001; Thompson et al., 2005; Seo et al., 2016). ATG8-PE is targeted to a pre-autophagosomal structure where it is thought to play a role in autophagosome formation. The clade II isoforms (ATG8h and ATG8i) lack extra amino acid residues at the C-terminus after the glycine residue, indicating that ATG8h and ATG8i proteins can interact with the autophagosome membrane without ATG4 processing (Seo et al., 2016).

Both pro-and anti-cell death functions have been linked to autophagy (Liu et al., 2005; Patel et al., 2008; Hofius et al., 2009; Yoshimoto et al., 2009; Wang et al., 2011; Munch et al., 2014; Xu et al., 2017). Silencing ATG6/Beclin1 leads to unrestricted spreading cell death beyond the sites of TMV infection on the leaves of *N. benthamiana* plants expressing the *N* gene (Liu et al., 2005), whereas HR triggered by RPS4 and RPM1 is compromised in *atg* mutants (Hofius et al., 2009). In addition, auto-immune phenotypes of chlorotic cell death/accelerated senescence is observed in different *atg* mutants (Xiong et al., 2005; Chung et al., 2006; Yoshimoto et al., 2009; Lai et al., 2011; Lenz et al., 2011; Wang et al., 2011). Depending on the lifestyles of pathogens, autophagy plays either a positive or a negative role in disease resistance (Hofius et al., 2017; Leary et al., 2019). It is now clear that autophagy plays a positive role in resistance to necrotrophic fungal pathogens (Lai et al., 2011; Lenz et al., 2011), but a negative role in SA-dependent resistance to biotrophic bacterial and fungal pathogens (Lai et al., 2011; Lenz et al., 2011; Wang et al., 2011).

Here, we showed that loss of function of CLC2 and CLC3 leads to chlorotic cell death/accelerated senescence in Arabidopsis resulting from elevated accumulation of both reactive oxygen species (ROS) and salicylic acid (SA). In addition, we provided genetic evidence that ROS-SA forms a positive feed-back loop that amplifies the autoimmune signals and concurrent presence of both ROS and SA is essential for the autoimmune phenotype displayed by *clc2-1clc3-1* plants. Unexpectedly, we found that loss of CLC2 and CLC3 impairs autophagy, which may account for the autoimmune phenotype of the *clc2-1clc3-1* mutant. Furthermore, we provided evidence that CLC2 directly interacted and co-localized with the Clade II ATG8 proteins, ATG8h and ATG8i, in an AIM1-and LDS-dependent manner. Lastly, we showed that both CLC2 and ATG8h/8i were subjected to autophagic degradation in the vacuoles. Taken together, our results uncovered a previously unidentified link between CLC function and autophagy in plants.

## Results

### Loss of the *Clathrin Light Chain 2* and *Clathrin Light Chain 3* in Arabidopsis results in SA-and H_2_O_2_-dependent activation of defense responses

While investigating the roles of the clathrin light chains in endocytosis of auxin efflux carriers (Wang et al., 2013; Zhang et al., 2017), it was unexpectedly found that accelerated senescence and/or spontaneous cell death occurred on the leaves of the *clc2-1clc3-1* double mutant plants (compare Figure 1A and 1B, see arrows in Figure 1B). The cell death was further confirmed by trypan blue staining (compare Figure 1C and 1D, see arrows in Figure 1D). Spontaneous cell death in plants is usually induced by over-accumulation of H_2_O_2_ (Lamb and Dixon, 1997). Consistent with this, the accumulation level of H_2_O_2_ was significantly higher on the leaves of the *clc2-1clc3-1* double mutant plants than on equivalent leaves of the wild-type Col-0 plants (compare Figure 1E and 1F, see arrows in Figure 1F).

**Figure 1.**
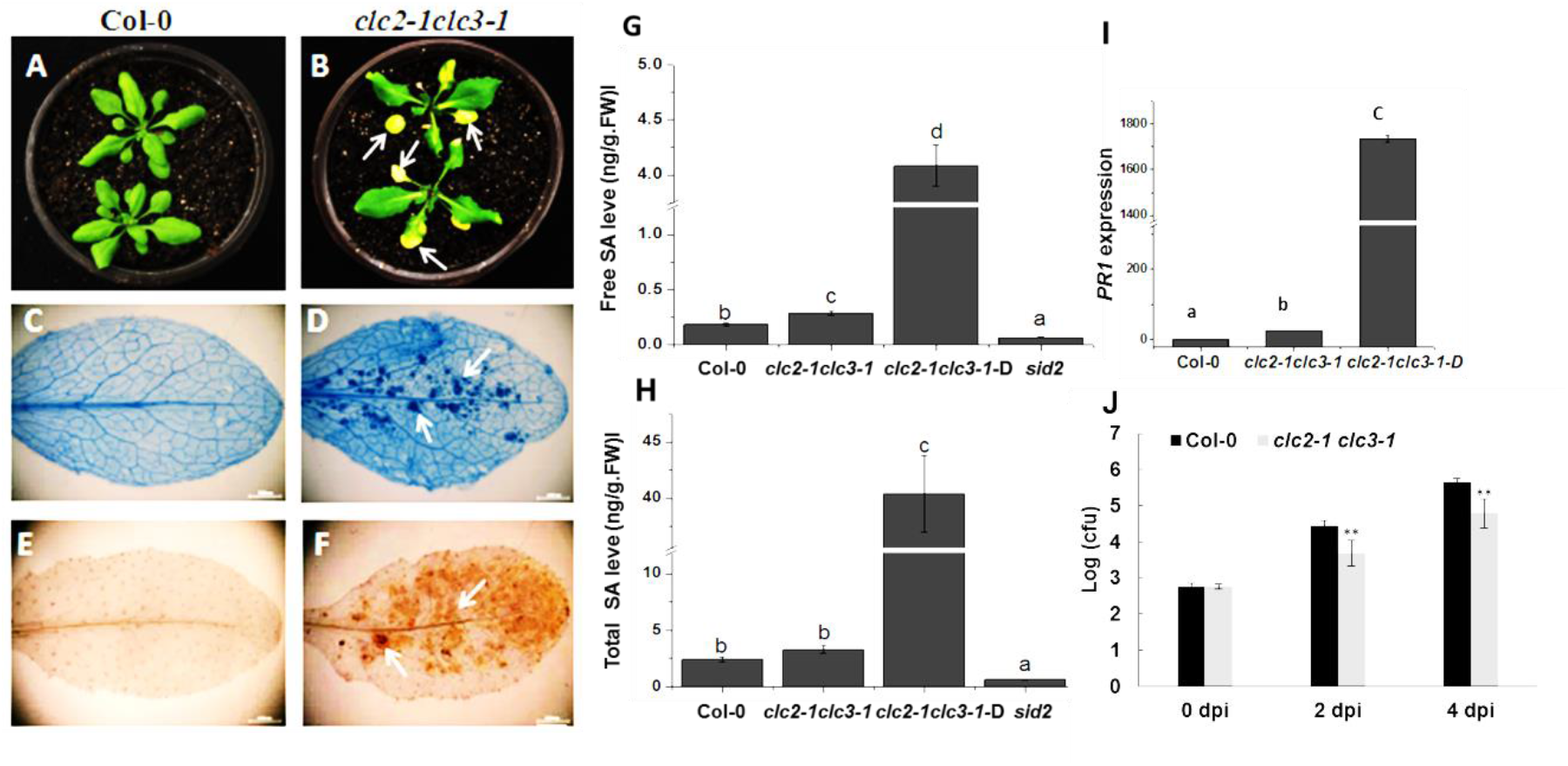
Defense responses are activated in the *clc2-1clc3-1* mutant. Wild-type Col-0 (**A**) and *clc2-1clc3-1* double mutant plants (**B**) were photographed at 30 days after sowing. Leaves of the *clc2-1clc3-1* double mutant plants exhibiting accelerated senescence or chlorotic cell death are indicated by the white arrows (**B**). Leaves of Col-0 (**C**) and *clc2-1clc3-1* double mutant plants (**D**) were stained with trypan blue. The white arrows in **D** point to examples of intense trypan blue staining indicative of cell death in the *clc2-1clc3-1* double mutant. Leaves of Col-0 (**E**) and *clc2-1clc3-1* double mutant plants (**F**) were stained with DAB to detect accumulation of H2O2. The white arrows in (**F**) point to examples of DAB staining indicative of H2O2 accumulation in the leaves of the *clc2-1clc3-1* double mutant. Elevated accumulation of free SA (**G**) and total SA (**H**) in the *clc2-1clc3-1* double mutant plants relative to Col-0 plants. (**I**) Expression of the *PR1* gene is highly induced in the *clc2-1clc3-1* double mutant relative to wild-type Col-0 plants. The *ACTIN2* (At3G18780) was used as the endogenous reference gene. The significant differences in (**G**), (**H**) and (**I**) were determined by an ANOVA with a post hoc Duncan’s test as indicated by different letters (P<0.05). (**J**) The *clc2-1clc3-1* double mutant plants displayed enhanced resistance against *Pseudomonas syringae pv. tomato* DC3000 (*Pst* DC3000). Asterisks indicate a significant difference from the control (**, *P* < 0.01, Student’s *t* test). The experiments were repeated three times with similar results.

Salicylic acid (SA) is a potent cell death inducer and it interacts with H_2_O_2_ synergistically to induce cell death (Shirasu et al., 1997; Yoshimoto et al., 2009; Xu et al., 2018). As expected, we found that both free SA and total SA levels were significantly higher in the leaves of *clc2-1clc3-1* double mutant plants than in that of Col-0 plants, especially in the leaves with cell death (Figure 1G and 1H). Together, these results indicate that the accelerated senescence and/or cell death observed in the *clc2-1clc3-1* plants is correlated with the over-accumulation of both H_2_O_2_ and SA.

Consistent with the enhanced H_2_O_2_ and SA levels, we found that expression of the *Pathogenesis-related gene 1* (*PR-1*) was induced to a much higher level in the *clc2-1clc3-1* mutant plants than in the Col-0 plants (Figure 1I). Accordingly, the *clc2-1clc3-1* mutant plants displayed an enhanced resistance against *Pseudomonas syringae pv.* DC3000 (*Pst* DC3000) relative to Col-0 plants (Figure 1J). These results indicate that loss function of CLC2 and CLC3 in Arabidopsis results in SA-and H_2_O_2_-associated cell death and activated defense responses.

### Loss function of either Isochorismate synthase 1 (ICS1) or Respiratory burst oxidase homolog protein D (RBOHD) rescues the autoimmune phenotypes of *clc2-1clc3-1* double mutant plants

*Salicylic acid induction deficient 2* (*SID2*) encodes the key enzyme, Isochorismate synthase 1 (ICS1), which converts isochorismate to chorismate in the chloroplast and is responsible for 90% pathogen-induced SA biosynthesis in Arabidopsis (Wildermuth et al., 2001). Respiratory burst oxidase homolog protein D (RBOHD) is the key subunit of the plasma-membrane-localized NAPDH oxidase, which is responsible for pathogen-induced H_2_O_2_ production (Torres et al., 2002). To genetically dissect the signaling pathways that are involved in the auto-immune phenotypes in the *clc2-1clc3-1* mutant, *clc2-1clc3-1sid2* and the *clc2-1clc3-1rbohD* triple mutants were generated by genetic crossing. Interestingly, deficiency of either *SID2* or *RBOHD* could rescue almost all the auto-immune phenotype displayed in the *clc2-1clc3-1* mutant plants, including the accelerated senescence or spontaneous cell death (Figure 2A and 2B), the enhanced accumulation of both H_2_O_2_ (Figure 2C) and SA (Figure 2D and 2E), the induced expression of *PR1* (Figure 2F), as well as the enhanced bacterial resistance (Figure 2G). These results strongly indicate that SA and ROS form a positive feedback loop that initiates and amplifies the immunity-related signals. The concurrent presence of both SA and H_2_O_2_ is required for the autoimmune responses observed in the *clc2-1clc3-1* double mutant plants.

**Figure 2.**
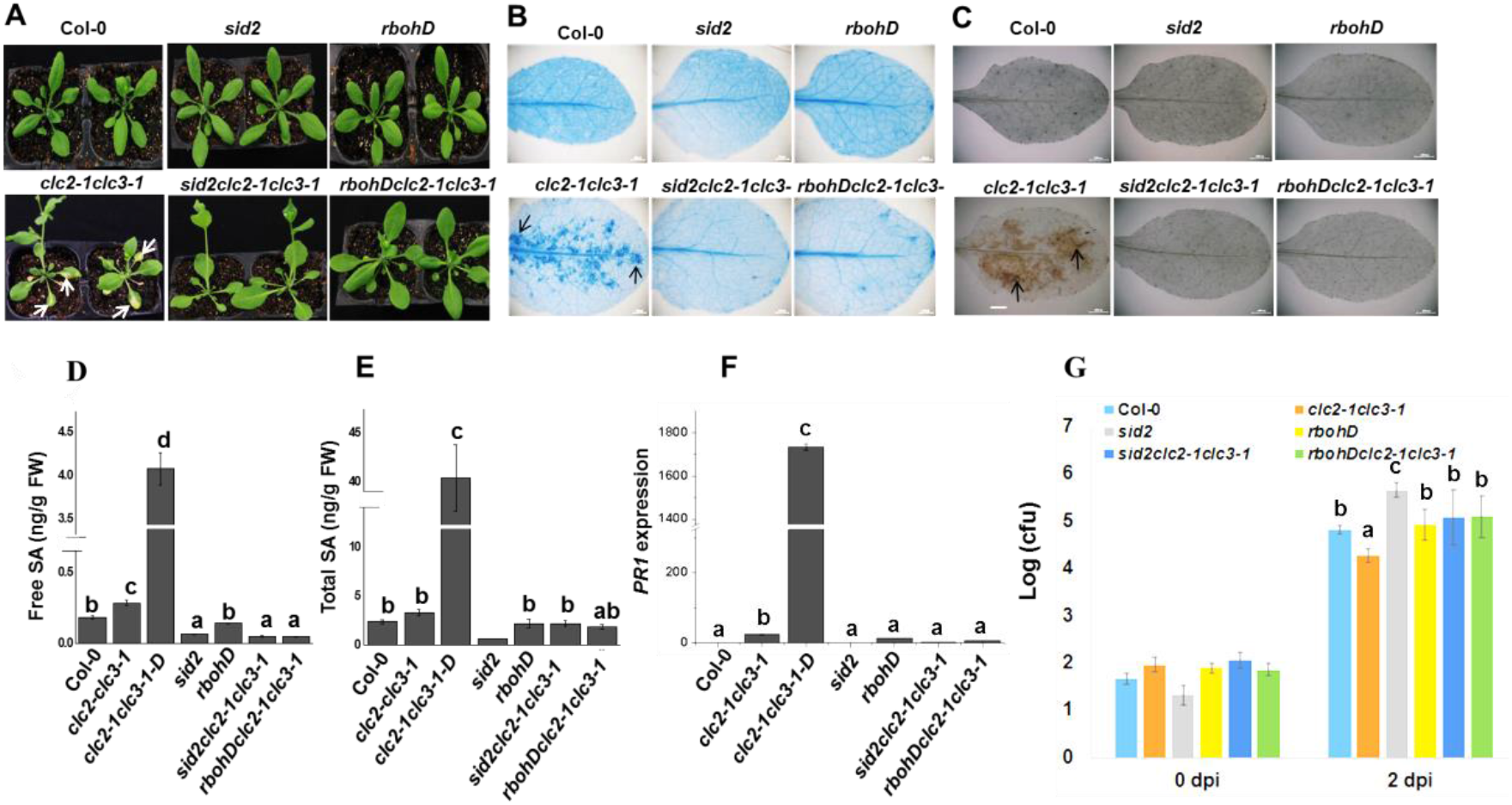
The auto-immune phenotype of the *clc2-1clc3-1* mutant is SA-and H2O2-dependent. Loss of function of either *ISC1*/*SID2* or *RBOHD* rescues the spontaneous defense responses observed in the *clc2-1clc3-1* mutant plants. (**A**) Wild-type Col-0 and mutant plants were photographed at 30 days after sowing. Leaves of the *clc2-1clc3-1* double mutant plants exhibiting accelerated senescence or chlorotic cell death are indicated by the white arrows. (**B**) Leaves of wild-type Col-0 and *clc2-1clc3-1* double mutant plants were stained with trypan blue. The white arrows point to examples of intense trypan blue staining indicative of cell death in the *clc2-1clc3-1* double mutant. (**C**) Leaves of wild-type Col-0 and mutant plants were stained with DAB to detect accumulation of H2O2. The white arrows in (**C**) point to examples of DAB staining indicative of H2O2 accumulation in the leaves of the *clc2-1clc3-1* double mutant. Levels of free SA (**D**), total SA (**E**), and expression of *PR1* mRNA (**F**) in wild-type Col-0 and mutant plants. The *ACTIN2* (At3G18780) was used as the endogenous reference gene. (**G**) Growth of *Pst* DC3000 in wild-type Col-0 and mutants at 0 and 2 days post-inoculation (dpi). The significant differences from (**D**) to **(G**) were determined by an ANOVA with a post hoc Duncan’s test as indicated by different letters (P<0.05).

### The *clc2-1clc3-1* double mutant phenocopies *atg2-1* mutant plants

The autoimmune phenotypes observed in the *clc2-1clc3-1* mutant (Figure 1) were also observed in many *atg* mutants (Wang et al., 2011; Lenz et al., 2011; Yoshimoto et al., 2009). The similarities shared between the *clc2-1clc3-1* and the *atg* mutants raise the possibility that clathrin function is somehow associated with autophagy pathway. To test this, we compared the phenotypes of the *atg2-1* mutant with that of the *clc2-1clc3-1* mutant side-by-side. We found that the *atg2-1* mutant plants displayed an accelerated senescence and cell death phenotypes similar to the *clc2-1clc3-1* (Figure S1A and S1B). In addition, H_2_O_2_ was similarly over-accumulated both in the *atg2-1* and the *clc2-1clc3-1* mutant plants (Figure S1C). Furthermore, it has been well documented that the autoimmune phenotype of many *atg* mutants is SA-dependent (Yoshimoto et al., 2009; Lenz et al., 2011; Wang et al., 2011) and SA highly accumulates in *atg* mutants relative to Col-0 (Yoshimoto et al., 2009; Lenz et al., 2011). Together, these results indicate that loss of CLC2 and CLC3 might have a negative impact on the autophagy pathway in Arabidopsis.

### clc2-1clc3-1 and atg2-1 mutant plants similarly display an enhanced sensitivity to carbon (C) and nitrogen (N) starvation

A hallmark of autophagy deficiency is enhanced sensitivity to C or N starvation (Li and Vierstra, 2012). To test whether the *clc2-1clc3-1* mutant is sensitive to N or C starvation, we performed N or C deprivation assays for seedlings of the indicated genotypes. The typical yellowish phenotype of enhanced sensitivity to either C or N starvation was observed for both the *clc2-1clc3-1* and the *atg2-1* seedlings but less so for the Col-0 seedlings (Figure 3A). In addition, we also treated the detached leaves from 4-week-old plants of indicated genotypes in the dark for 6 days. Consistent with the C-deprivation experiments with seedlings, an accelerated senescence was also observed for the dark-treated *clc2-1clc3-1* and *atg2-1* mutant leaves but not for the dark-treated Col-0 leaves (Figure 3B). The chlorophyll contents measured were highly correlated with the chlorotic phenotypes among these lines (Figure 3C). Collectively, these data strongly indicated that autophagy is likely compromised in the *clc2-1clc3-1* mutant.

**Figure 3.**
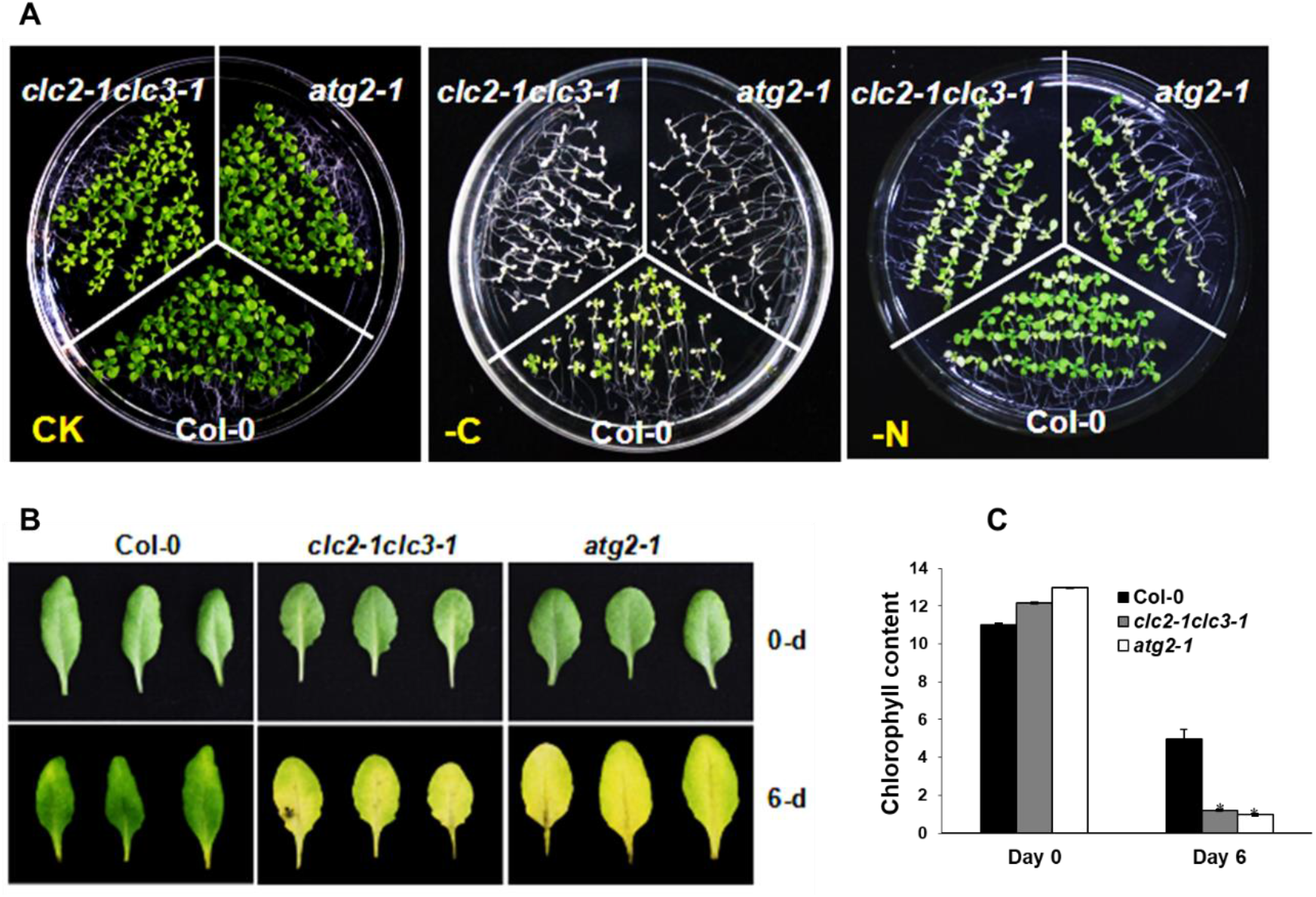
The *clc2-1clc3-1* mutant exhibits enhanced sensitivity to carbon and nitrogen starvation. **(A**) Phenotype of the *clc2-1clc3-1* double mutant under C and N starvation condition. Seeds were germinated and grown on 1/2 MS medium plates supplemented with both C and N (+C+N) and without C (-C) or N (-N) under normal growth conditions. At 7 days post germination, the +C and-N seedlings were kept under normal growth conditions and the-C seedlings were placed in the dark for accelerated C deficiency. After dark treatment for 10 days, the-C seedlings were allowed to recover under normal conditions for additional 10 days, while the CK and-N seedlings were grown under normal conditions during the entire process. (**B**) Phenotype of the dark-treated leaves from the indicated genotypes. Detached leaves from the 4-week-old plants were put onto a piece of wet filter paper in petri dishes and wrapped with aluminum foil for dark treatment. The images were taken at 0 day (upper panel) and 6 days post dark treatment (lower panel). (**C**) The relative chlorophyll content of the Col-0, *clc2-1clc3-1* and *atg2-1* seedlings. The relative chlorophyll content was calculated by comparing the values in-C or-N seedlings versus +C or +N seedlings. The experiments were repeated 3 times with similar results.

### The *clc2-1clc3-1* mutant plants exhibit enhanced resistance against a bacterial pathogen *Pseudomonas syringae pv.* tomato DC 3000 and a fungal pathogen *Golovinomyces cichoracearum*

It has been reported that the *atg2-1* mutant plants display an enhanced resistance against a biotrophic fungal pathogen, *Golovinomyces cichoracearum,* UCSC1 (Wang et al., 2011). We therefore performed resistance assays to compare the *G. cichoracearum* resistance of the *clc2-1clc3-1* mutant with that of wild-type Col-0 and the *atg2-1* mutant. At 7 days post inoculation (dpi), aniline blue staining was performed to monitor cell death, production of conidiophores, and growth of hyphae on the infected leaves. Cell death developed on the infected leaves of both the *clc2-1clc3-1* and the *atg2-1* mutant plants but not on the infected Col-0 leaves (Figure 4A, see arrow-pointed regions). In addition, abundant fungal hyphae and conidiophores were visible on the leaves of the Col-0 plants, whereas much fewer hyphae and conidiophores were seen on the leaves of the *clc2-1clc3-1* and the *atg2-1* mutant plants (Figure 4B). To accurately measure fungal reproduction, the conidiophores were counted on the infected leaves at 7 dpi. As shown in Figure 4C, the number of conidiophores formed on the leaves of the *clc2-1clc3-1* mutant plants was significantly less than on the Col-0 plants, indicating that the *G. cichoracearum* resistance is significantly enhanced in the *clc2-1clc3-1* mutant plants.

**Figure 4.**
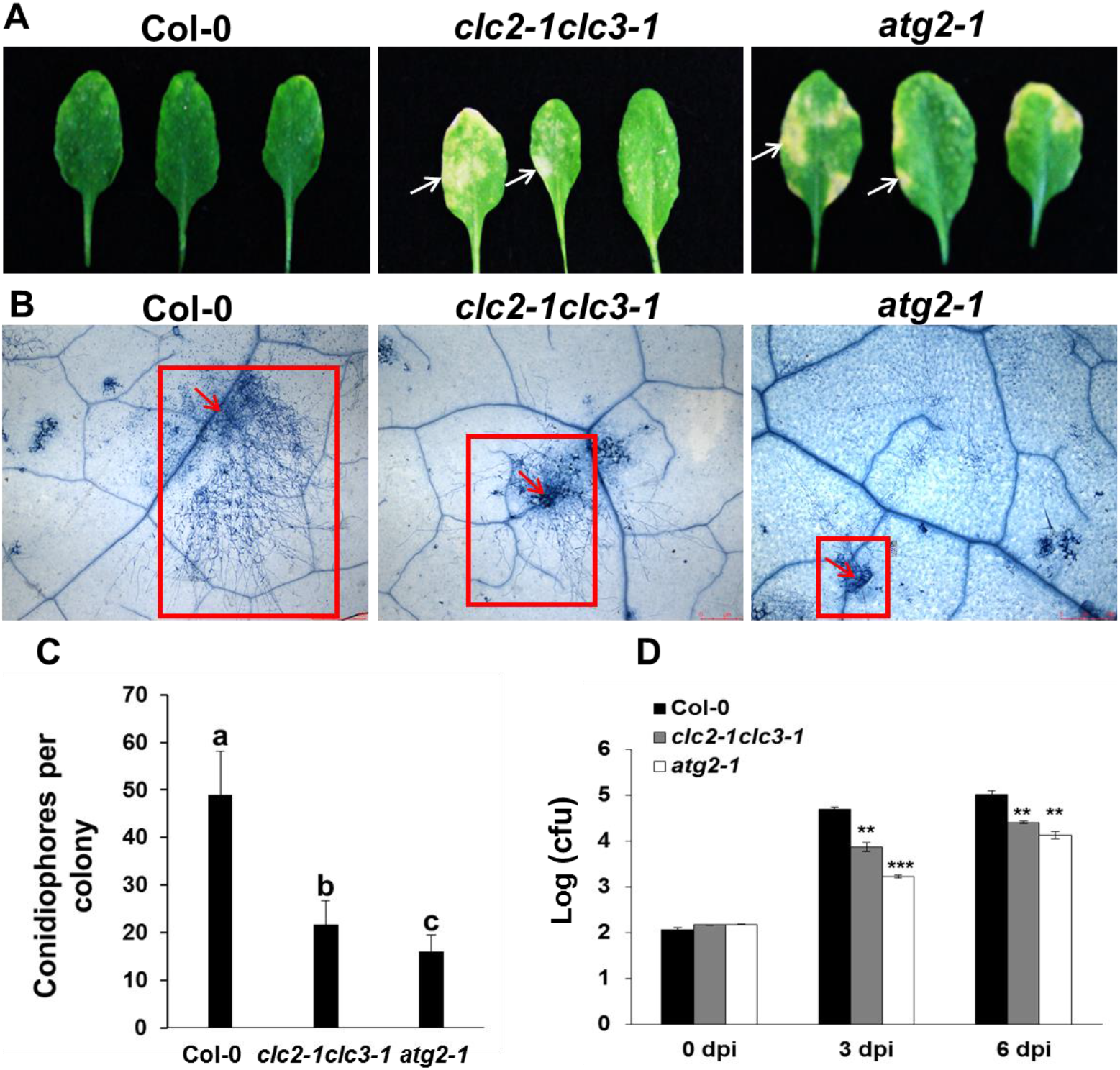
The *clc2-1clc3-1* mutant exhibits enhanced resistance to *Golovinomyces cichoracearum* and *Pst* DC3000. **(A**) Comparison of the symptoms of 6-week-old Col-0, *clc2-1clc3-1*, and *atg2-1* plants grown under short day condition (8h-light/16h-dark) infected with *G. cichoracearum*. The photos were taken at 7 days post-inoculation. The cell death caused by infection is indicated by arrows. (**B**) Conidiophores and hyphal growth of *G. cichoracearum* on the infected leaves shown in (**A**) were visualized by Trypan blue staining. The boundary of hyphae derived from a single colony is indicated by the red rectangles. (**C**) The number of conidiophores formed on the leaves of the different genotypes. Bars represent mean and standard deviation of one representative experiment. ** indicates statistically significant differences (P < 0.01; Student’s t-test, n=22). (**D**) The *clc2-1clc3-1* double mutant and the *atg2-1* mutant plants displayed enhanced resistance against *Pst* DC3000. cfu represents colony forming unit. Asterisks indicate a significant difference from the control (** represents *P* < 0.01, and *** represents P < 0.001, Student’s *t* test). The experiments were repeated 3 times with similar results.

We have shown that the *clc2-1clc3-1* mutant plants displayed enhanced resistance against the biotrophic bacterial pathogen *Pst* DC3000 (Figure 1J). However, the resistance of the *atg* mutants to *Pst* DC3000 remains controversial (Hofius et al., 2009; Lenz et al., 2011; Wang et al., 2011). To resolve this issue, we performed bacterial growth assays. As shown in Figure 4D, the growth of the *Pst* DC3000 was significantly lower on the leaves of both the *clc2-1clc3-1* and the *atg2-1* mutant plants than on the leaves of the Col-0 plants, indicating that *clc2-1clc3-1* and *atg2-1* mutant plants exhibit enhanced resistance against *Pst* DC3000. Together, our results indicate that the *clc2-1clc3-1* and the *atg2-1* mutants display similar resistance against two different types of biotrophic pathogens.

### Autophagy is compromised in the *clc2-1clc3-1* mutant plants

The similarities shared between the *clc2-1clc3-1* and the *atg2-1* mutant plants (Figure 3, Figure 4 and Figure 1S) raise the possibility that autophagy is compromised in the *clc2-1clc3-1* mutant. Various assays to monitor autophagy in plants have been developed (Li and Vierstra, 2012). The fusion proteins between ATG8s and fluorescence proteins (FPs) have been used as a marker to visualize autophagosomes or autophagic bodies *in vivo* (Yoshimoto et al., 2004; Yoshimoto et al., 2009; Wang et al., 2011). In Arabidopsis, ATG8-labeled vesicles fail to accumulate in autophagy mutants (Chung et al., 2010; Wang et al., 2011). To examine whether the *clc2-1clc3-1* mutant exhibits autophagy defects, accumulation of autophagic bodies represented by GFP-ATG8e transgenically expressed in the roots of 5-day-old Col-0 and *clc2-1clc3-1* seedlings was examined. A significantly reduced number of the autophagic bodies was observed in the root cells of *clc2-1clc3-1* mutant seedlings compared to that of the Col-0 seedlings (Figure 5A and 5B), indicating that the autophagic pathway was compromised in the *clc2-1clc3-1* mutant.

**Fig. 5.**
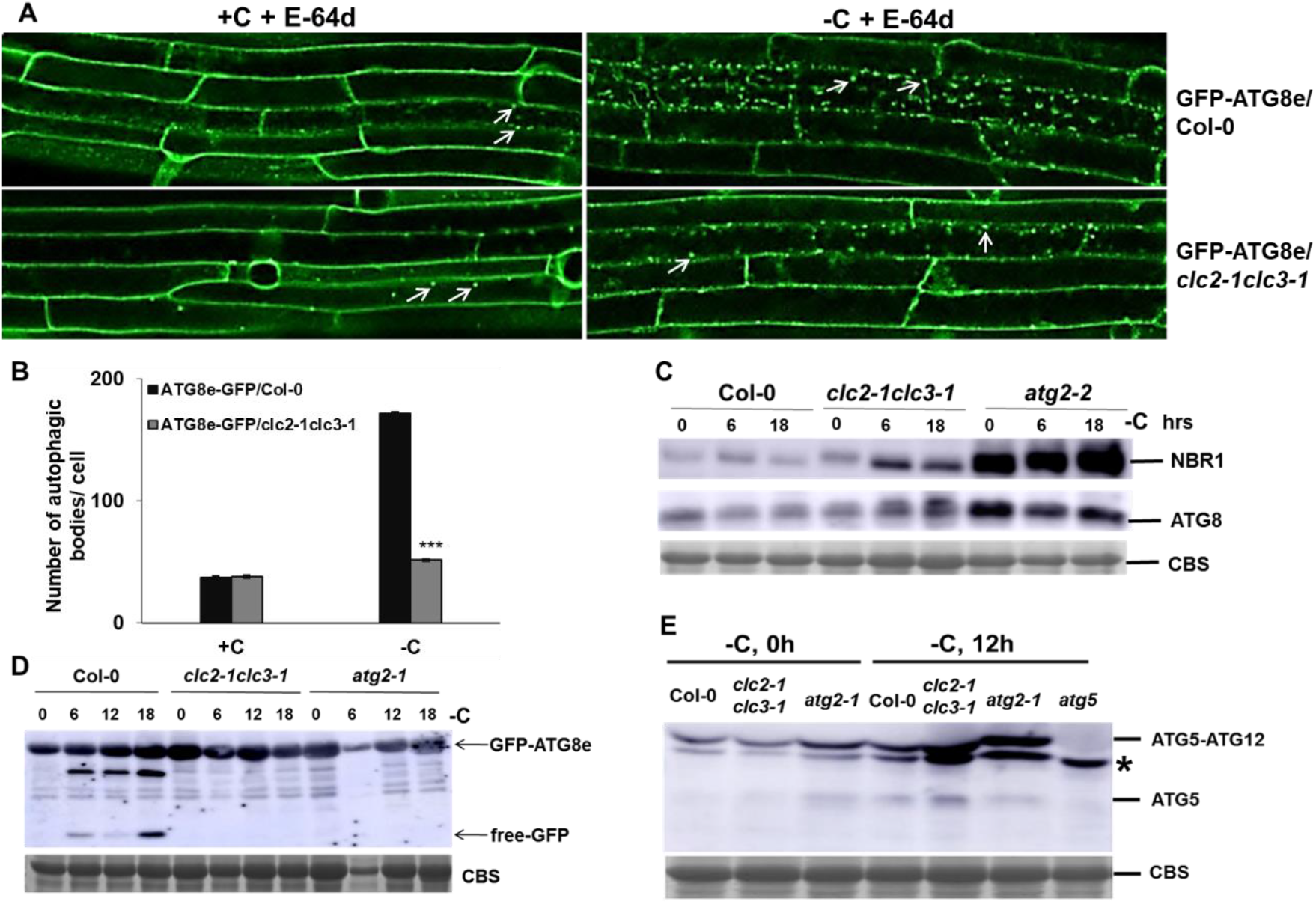
Autophagy is compromised in the *clc2-1clc3-1* mutant. (**A**) The number of autophagic bodies tagged by the transgenically expressed GFP-ATG8e was significantly reduced in the *clc2-1clc3-1* mutant seedlings. Five-day-old seedlings grown in 1/2 MS medium containing sucrose (+C, **left panel**) or no sucrose (-C, **righ**t **panel**) were treated with E64-d, a protease inhibitor, for 2 hours. Primary root cells were observed and the images of the GFP-ATG8e bodies were captured by confocal microscopy. (**B**) Quantification of the GFP-ATG8e bodies shown in panel A (n>10). (**C**) Autophagic degradation of both NBR1 (**top panel**) and ATG8 (**middle panel**) was attenuated in the *clc2-1clc3-1* mutant leaves. Coomassie blue staining (CBS) was used as a loading control (**lower panel**). Proteins were extracted from the detached leaves of 4-week-old plants of the indicated genotypes that were dark-treated for different periods of time as labeled. Western blotting was performed using anti-AtNBR1 and anti-AtATG8, respectively. (**D**) Free GFP was released from GFP–ATG8e in WT but not in the *clc2-1clc3-1* and the *atg2-2* seedlings exposed to C-deficient medium. Proteins were extracted from the detached leaves of 4-week-old plants of the indicated genotypes that were dark-treated for different periods of time as labeled. The Western blotting was performed using a GFP-specific antibody. CBS was used as a loading control (lower panel). (**E**) Immunoblot analysis of the ATG12-ATG5 conjugate in plants of the indicated genotypes with an Arabidopsis ATG5-specific antibody under normal (+C) and C starvation (-C) conditions. Equal protein loading was confirmed by CBS staining. A non-specific band was indicated by *. The experiment was repeated 3 times with similar result.

Accumulation of both ATG8s, the structural components of autophagosomes and NBR1, a selective receptor for ubiquitinated proteins, indicates a defect in the autophagy pathway (Yoshimoto et al., 2004; Thompson et al., 2005; Thompson and Vierstra, 2005; Phillips et al., 2008). To further confirm that the autophagy pathway is indeed impaired in the *clc2-1clc3-1* mutant, we monitored the abundance of ATG8 and NBR1. As shown in Figure 5C, under C starvation condition, a moderate increased accumulation of both ATG8 and NBR1 was observed in the *clc2-1clc3-1* mutant plants relative to the Col-0 plants, confirming that both bulk autophagy and NBR1-mediated selective autophagy are compromised in *clc2-1clc3-1* mutant. As a control, a substantially enhanced accumulation of both ATG8 and NBR1 was observed in the *atg2-1* mutant even under normal conditions (Figure 5C). The accumulation level of both ATG8 and NBR1 was higher in the *atg2-1* mutant than in the *clc2-1clc3-1* mutant regardless of the C status (Figure 5C).

The vacuolar accumulation of free GFP, which is released from the GFP-ATG8 fusions during breakdown of the autophagic bodies measures autophagy-dependent vacuolar transport, and the free GFP accumulation in the vacuole is almost completely absent in mutants that block ATG8 lipidation (Yoshimoto et al., 2004; Chung et al., 2010; Suttangkakul et al., 2011). Thus, we compared the accumulation of vacuolar free GFP between Col-0, *clc2-1clc3-1*, and *atg2-2* plants transgenically expressing ATG8e-GFP by Western blotting. As shown in Figure 5D, the release of free GFP was only seen in the dark-treated ATG8e-GFP/Col-0 plants but not in either the ATG8e-GFP/*clc2-1clc3-1* or the ATG8e-GFP/*atg2-2* plants, further confirming that autophagy is impaired in the *clc2-1clc3-1* mutant.

### The accumulation level of the ATG12-ATG5 complex is elevated in the clc2-1clc3-1 mutant

The formation of the ATG12–ATG5 protein complex is essential for ATG8 lipidation (Chung et al., 2010). In addition to ATG5, the ATG12–ATG5 protein complex can also be detected by the anti-ATG5 on a Western blot (Chung et al., 2010). To examine whether loss function of CLC2 and CLC3 interferes with the formation of the ATG12– ATG5 conjugate, Western blot analysis using an Arabidopsis ATG5-specific antibody was performed on the protein samples extracted from the leaves of 4-week-old plants either treated or not treated under dark for 12 hours. As shown in Figure 5E, under both light and dark conditions, the formation of the ATG12–ATG5 protein complex was observed in the *clc2-1clc3-1* mutant, indicating loss function of CLC2 and CLC3 does not interfere with the formation of the ATG12–ATG5 complex. Interestingly, we found that the formation of the ATG12–ATG5 complex was also observed in the *atg2-1* mutant, consistent with the fact that ATG2 functions in a distinct step of autophagy unrelated to ATG5 and ATG12 (Li and Vierstra, 2012). To our surprise, we found that the formation of the ATG12–ATG5 complex was actually increased to a much higher level in both the *clc2-1clc3-1* and the *atg2-1* mutant plants relative in Col-0 plants under dark condition (Figure 5E). A similar increase in accumulation of ATG12-ATG5 was also observed in other *atg* mutants, which is consistent with idea that blocking autophagy up-regulates expression of *ATG* genes and suppresses the turnover of ATG proteins (Thompson et al., 2005; Phillips et al., 2008; Chung et al., 2010; Li et al., 2014; Huang et al., 2019; Liu et al., 2020). This result suggests again that autophagy is compromised in the *clc2-1clc3-1* mutant. As expected, neither ATG5 nor the ATG12-ATG5 complex was detectable in the *atg5* mutant (Figure 5E).

### Enhanced level of ROS was accumulated in the chloroplasts of clc2-1clc3-1 mutant plants relative to Col-0

Chloroplasts are one of the major sources of secondary ROS production (Mittler, 2017). Autophagy plays a critical role in maintaining cellular ROS homeostasis by reducing secondary ROS production through eliminating damaged chloroplasts (Ishida et al., 2008; Izumi et al., 2010; Xie et al., 2015). We reasoned that the enhanced accumulation of H_2_O_2_ in the *clc2-1clc3-1* mutant (Figure 1F) could be due to over-accumulation of dysfunctional chloroplasts as a result of impaired autophagy pathway. To test this possibility, H_2_DCFDA staining was performed to compare the ROS accumulation in the chloroplasts of Col-0 and *clc2-1clc3-1* mutant plants treated under dark. As shown in Figure 6, the ROS level was significantly higher in the chloroplasts of *clc2-1clc3-1* and *atg2-1* mutant plants than in Col-0, suggesting that compromised chlorophagy in *clc2-1clc3-1* mutant leads to over-accumulation of damaged chloroplasts, in which secondary ROS is produced.

**Figure 6.**
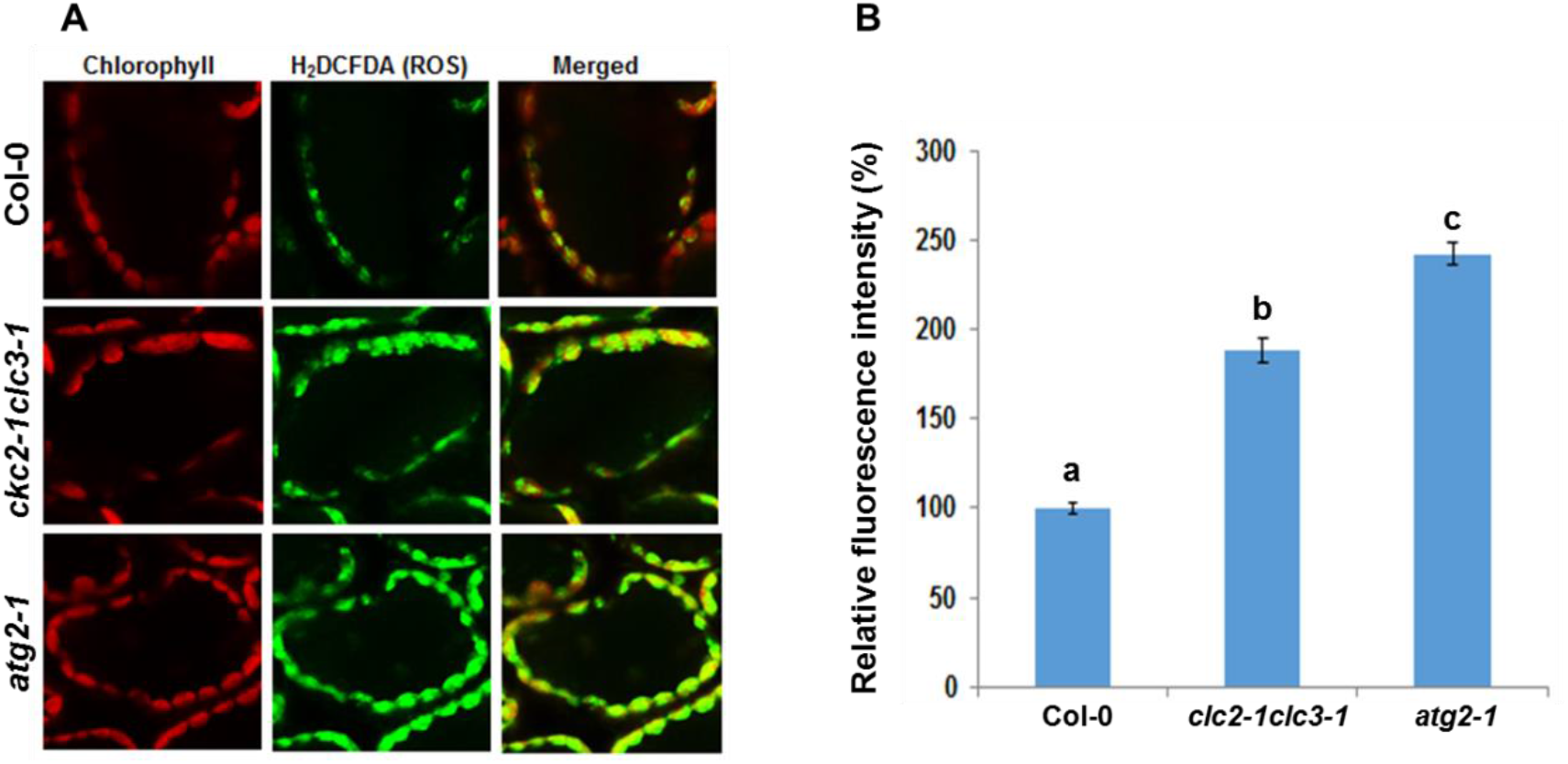
ROS accumulation is elevated in the chloroplasts of mesophyll cells of *clc2-1clc3-1* seedlings relative to Col-0. (**A**) The presence of ROS in the chloroplasts of mesophyll cells of the indicated genotypes was visualized by staining with H2DCFDA. Auto-fluorescence of chlorophyll (red, left) and ROS fluorescence (green, middle) were captured by a confocal microscopy. The merged images of red and green channels were shown on the right; (**B**) Quantification of the green fluorescence signals shown in (**A**). The intensities of the green fluorescence signals, which represent ROS levels, were quantified for at least 20 cells of each genotype by ImageJ. The data are shown as means±SE (*n*>20). Different letters represent significant difference at 0.05 using ANOVA with a posthoc Duncan’s test. The experiment was repeated twice with similar results.

### CLC2 but not CLC3 directly interacts with ATG8h and ATG8i

A CLC subunit was identified as an ATG8-interacting protein in *N. benthamiana* (Macharia et al., 2019), raising a possibility that CLCs might directly interact with ATG8s. To test the possibility, we performed Yeast-2-hybrid (Y2H) assays of CLC2 or CLC3 with eight out of nine ATG8 variants (ATG8a-ATG8i) except ATG8b in Arabidopsis. To our surprise, we found that AD-CLC2 and not AD-CLC3 strongly interacted with BD-ATG8h and BD-ATG8i (Figure 7A and Figure 2S), but only weakly interacted with ATG8a, ATG8c, ATG8d and ATG8f when incubation time was extended (Figure 3S), suggesting that CLC2 participates in autophagy mainly through direct interaction with ATG8h and ATG8i. Coincidently, ATG8h and ATG8i are clustered into a closely related sub-clade (Figure 4S). The clade II isoforms (ATG8h and ATG8i) lack extra amino acid residues at the C-terminus after the glycine residue, implying that ATG8h and ATG8i proteins can interact with the autophagosome membrane without ATG4 processing (Figure 5S; Seo et al., 2016).

**Fig. 7.**
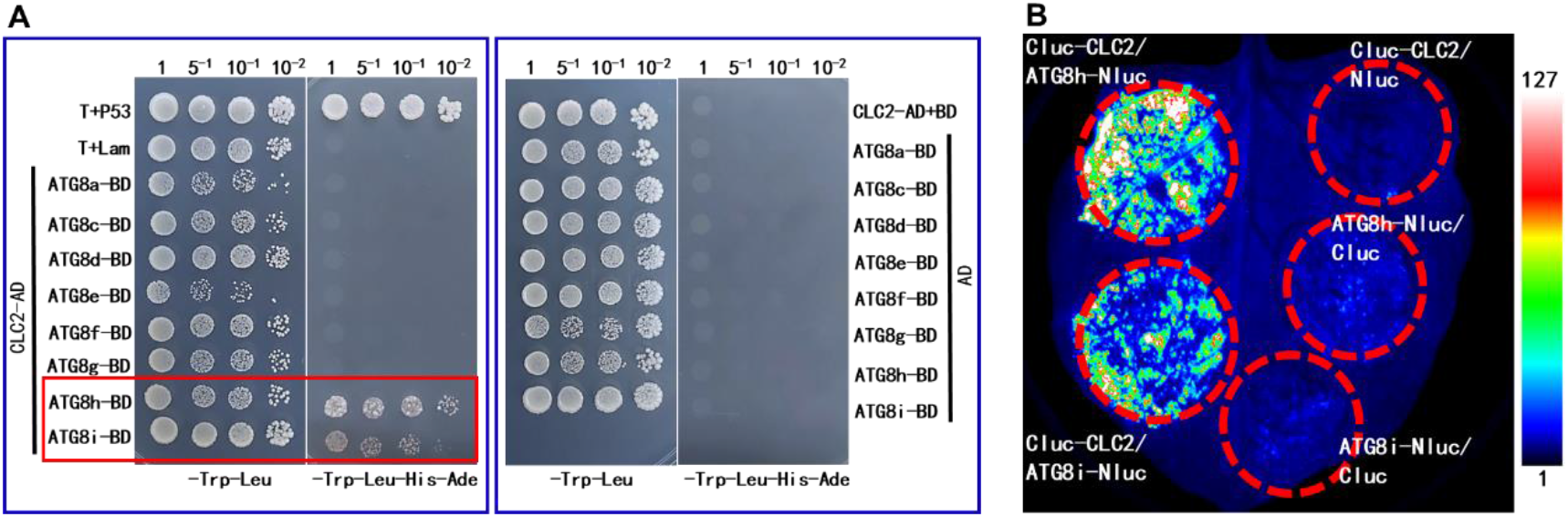
CLC2 interacts with ATG8h and ATG8i. (**A**) CLC2 interacts specifically with ATG8h-BD and ATG8i-BD but not with the other ATG8s and the empty BD vector in Y2H assays. Yeast cells on the left were selected on the SD/Trp^-^Leu^-^ medium and yeast cell on the right were selected on the SD/Trp^-^Leu^-^His^-^Ade^-^ medium; (**B**) cLuc-CLC2 interacts with ATG8h-nLuc and ATG8i-nLuc in a luciferase complementation assay.

To further confirm the CLC2-ATG8h or CLC2-ATG8i interaction, the ORFs of respective genes were cloned into binary Luciferase Complementation Imaging vectors (Chen et al., 2008). The constructed plasmids were then transiently co-expressed on the leaves of *N. benthamiana* via agro-infiltration. Consistent with the Y2H results (Figure 7A), luciferase activity was only observed when cLuc-CLC2 was co-infiltrated with ATG8h-nLuc or ATG8i-nLuc (Fig. 7B). No luciferase activity was detectable when cLuc-CLC2 co-infiltrated with nLuc or cLuc co-infiltrated with either ATG8h-nLuc or ATG8i-nLuc (Figure 7B). These results confirmed that CLC2 indeed interacts with ATG8h and ATG8i.

### CLC2 interacts with ATG8h through a unique ATG8-interacting motif (AIM) that not present in CLC3 and the LIR/AIM-docking site (LDS) presented in ATG8h

It has been established that cargo receptors interact with ATG8 proteins anchored on the autophagosome membrane through AIM for cargo recruitment (Noda et al., 2010). Two putative AIMs were identified in the amino acid sequence of CLC2 and CLC3, respectively (Figure 8A and Figure 6S). In addition to a common AIM shared between CLC2 and CLC3, both CLC2 and CLC3 have a unique AIM (Figure 6S). The fact that CLC3 did no interact with ATG8h/8i (Figure 2S) strongly suggests that the CLC2-ATG8h/8i interaction is most likely mediated by the AIM1 that is unique to CLC2 near its N terminus (Figure 8A and Figure 6S). To explore this possibility, an AIM1 deletion version (△AIM1) of CLC2 as well as a version with first and fourth amino acid of the AIM2 (**<u>W</u>**KA**<u>I</u>**) mutated to Ala (**<u>A</u>**KA**<u>A</u>**) were generated respectively, by site-directed mutagenesis, and tested for their interactions with ATG8h using both Y2H and Luciferase complementation assays. Consistent with our postulation, only deletion of AIM1 but not mutations at AIM2 abolished the CLC2-ATG8h interaction (Figure 8B and 8C), indicating that CLC2 interacts with ATG8h in an AIM1-dependent manner.

**Fig. 8.**
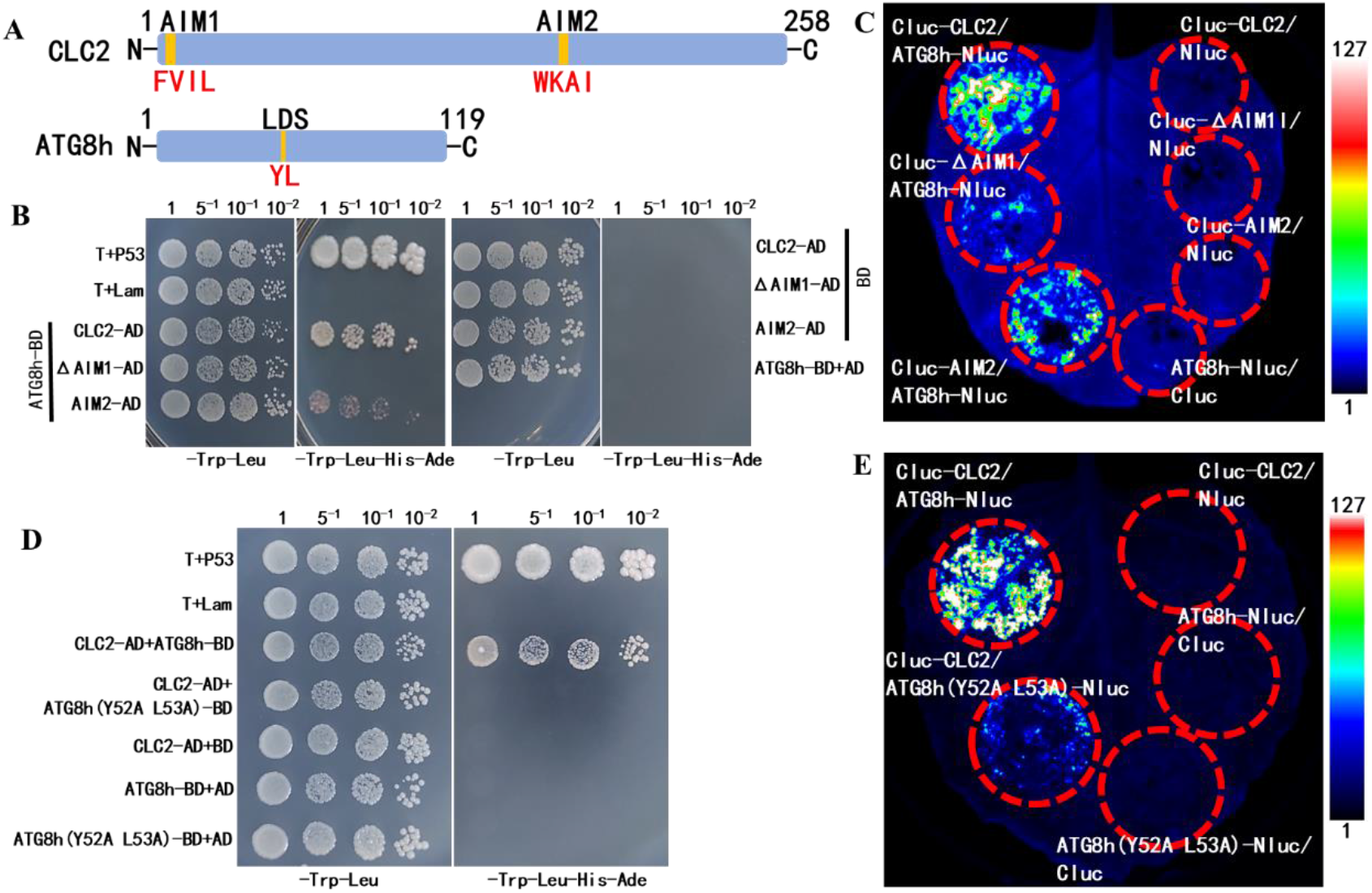
CLC2 interacts with ATG8h and ATG8i in an AIM1-and LDS-dependent manner. (**A**) The diagrams showing the locations of AIMs and LDS in CLC2 and ATG8h, respectively; (**B**) The interaction between CLC2 and ATG8h was abolished only by AIM1 deletion (△AIM1) but not by AIM2 mutations in a Y2H assay; (**C**) The luminescence reflecting the interaction between cLuc-CLC2 and ATG8h-nLuc was abolished only by AIM1 deletion (△AIM1) but not by AIM2 mutations; (**D**) The interaction between CLC2 and ATG8h was abolished by LDS mutations at Y52A, R53A within ATG8h in a Y2H assay; (**E**) The interaction between CLC2 and ATG8h was abolished by LDS mutations at Y52A, R53A within ATG8h in a luciferase complementation assay. In (**A**), (**C**), and (**E**), T-antigen and p53 were included as a positive interaction control, and Lam and T-antigen were included as a negative interaction control; The photos shown in Fig. **8A**, **8C** and **8E** were taken at 4 days post incubation. AD: activation domain; BD: binding domain. The numbers on top of the images are the diluting factors. In (**B**), (**D**) and (**F**), the luciferase complementation assay was performed by agro-infiltration of the indicated pairs of constructs into *N. benthamina* leaves. The images were captured by a CCD imaging system at 2 days post infiltration (dpi). These experiments were repeated 3 times with similar results.

The LIR/AIM docking site (LDS) presented in the ATG8 protein family is critical for the interactions between ATG8 proteins and autophagic receptors/adaptors, which leads to the recruitment of cargos to autophagosomes. A hydrophobic pocket is highly conserved in the Arabidopsis ATG8 isoforms in the LDS (Figure 8A and Figure 5S; Marshall et al., 2019; and Sun et al., 2022,). Mutations of Y50 and L51 simultaneously at the hydrophobic pocket to Alanine (A) (Y50A, L51A) abolished the LDS binding affinity of ATG8a or ATG8f (Marshall et al., 2019; and Sun et al., 2022). Therefore, we generated Y52A/L53A mutant of ATG8h by site-directed mutagenesis. Consistent with previous reports, we showed that Y52A/L53A substitutions in the ATG8h significantly weakened the interaction with CLC2 in the Y2H assay (Figure 8D), as well as in the split luciferase assay (Figure 8E). Together, these results indicated that both AIM1 in the CLC2 and LDS in the ATG8h are essential for CLC2-ATG8h

### Both GFP-ATG8h/GFP-ATG8i and CLC2-MBP are subjected to autophagic degradation

ATG8 proteins decorate the membranes of autophagosomes usually destine in the vacuoles for autophagic degradation. To test whether ATG8h and ATG8i similarly enter vacuoles and are subjected to autophagic degradation, transgenic seedlings of the *35S::GFP-ATG8h* and *35S::GFP-ATG8i* were generated and the effects of the dark treatment as well as dark + ConA, an inhibitor of autophagic degradation, were examined on the degradation of the autophagic bodies labelled by GFP in the root cells. Under the long-day conditions (16h-light/8h-dark), only a few autophagic bodies labelled by GFP-ATG8h or GFP-ATG8i were observed (Figure 9A and 9B, upper panels, see white arrows). Under autophagy induced conditions (constant dark treatment for 48-h), a defused form of fluorescence was observed in the root cells of both the *35S::GFP-ATG8h* and *35S::GFP-ATG8i* seedlings (Figure 9A and 9B, middle panels), indicating that the autophagic bodies labelled by GFP-ATG8h or GFP-ATG8i were mostly degraded into free GFP inside the vacuoles. However, upon dark treatment followed by ConA treatment, numerous autophagic bodies labelled by GFP-ATG8h or GFP-ATG8i were observed as a result of inhibition of autophagic degradation (Figure 9A and 9B, lower panels, see white arrows). Together, these results demonstrated that, like other ATG8 proteins, both GFP-ATG8h and GFP-ATG8i enter the vacuoles and are subjected to autophagic degradation.

**Figure 9.**
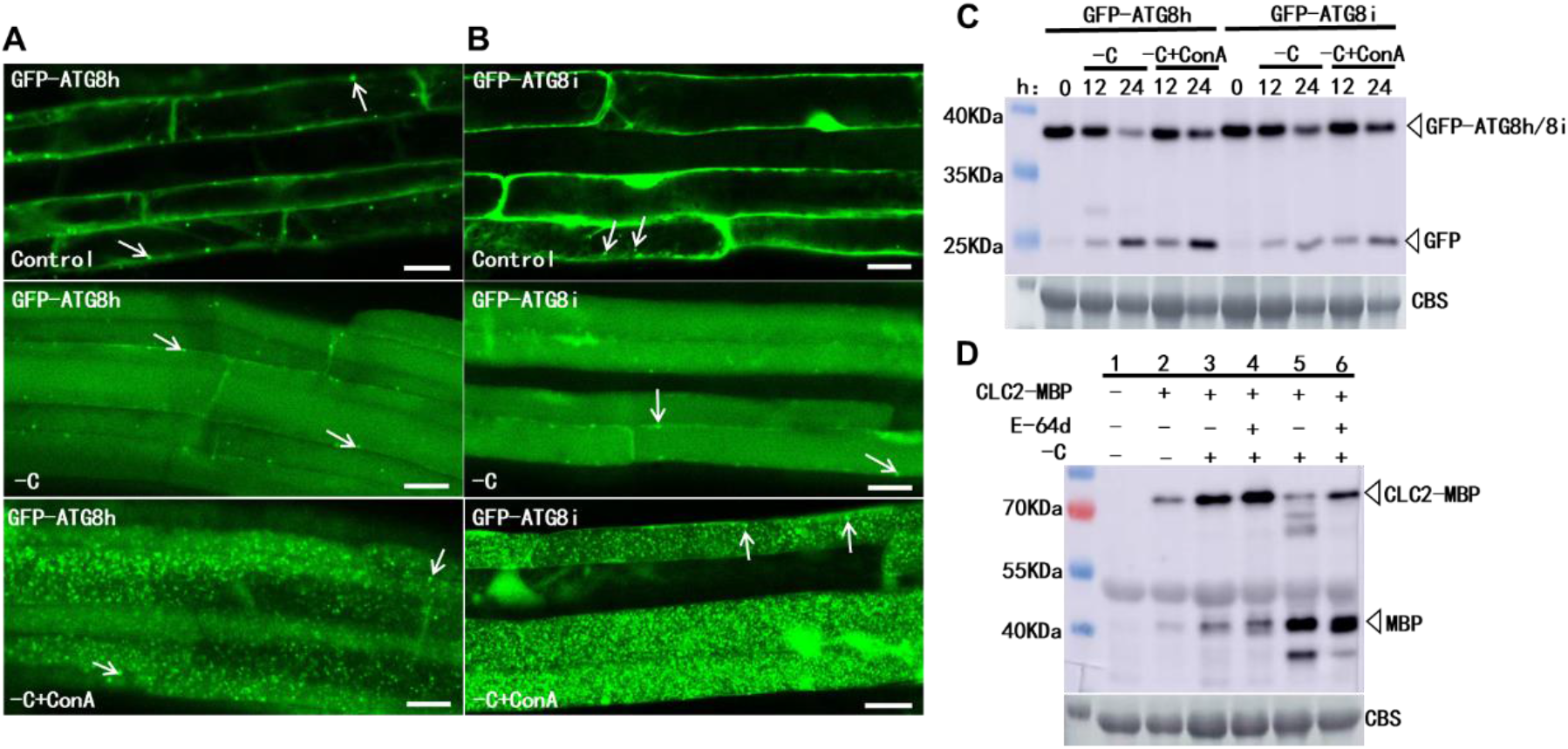
Detection of the autophagic degradations of GFP-ATG8h, GFP-ATG8i and CLC2-MBP. (**A and B**) Seven-day-old Col-0 seedlings expressing GFP-ATG8h or GFP-ATG8i were grown under C-sufficient conditions for 48-h (**upper panel**); under C-deprivation conditions for 48-h (**middle panel**); or under C-deprivation conditions for 36-h followed by adding 1 µM ConA for additional 12-h (**lower panel**). The GFP signals in the root cells were visualized by a laser confocal microscopy. White arrows pointed to the autophagosome or autophagic body; (**C**) Free GFP was released from GFP–ATG8h and GFP-ATG8i under autophagy inducing conditions. Seven-day-old GFP-ATG8h/WT and GFP-ATG8i/WT seedlings were grown in C-deficient liquid medium with or without containing 1 µM ConA for 12-h and 24-h respectively. The total protein was extracted from the respective seedlings and the free GFP released from GFP-ATG8h or GFP-ATG8i as a result of autophagic degradation was detected by Western blotting analysis with GFP antibody. CBS was used as a loading control. (**D**) Free MBP was released from CLC2-MBP under autophagy inducing conditions. The Agrobacterium strain GV3101 carrying *35S::CLC2-MBP* construct was infiltrated into the leaves of the *Nicotiana benthamiana* plants. Protein sample extracted from the leaves without infiltration of *35S::CLC2-MBP* (**Lane 1**); Protein sample extracted from the infiltrated leaves at 48-h post infiltration of *35S::CLC2-MBP* (**Lane 2**); The infiltrated leaves were cut at the base of petioles at 48-h post infiltration of *35S::CLC2-MBP*. The whole leaves were transferred into a-C liquid medium and incubated for additional 12-h (with the petioles in the medium, **Lane 3**); Similar to Lane 3, except that the whole leaves were transferred into a-C liquid medium that containing 100 mM E-64d and incubated for 12-h (**Lane 4**); The infiltrated leaf area was cut into 1 cm2 pieces and incubated in a C-deficient medium under dark for 36-h. These leaf discs were then transferred to C-deficient medium without containing 100 mM E-64d (**Lane 5**) or containing 100 mM E-64d (**Lane 6**) in the dark for additional 12-h. The total proteins were extracted from leaf samples treated under different conditions (see description above), and the free MBP released from CLC2-MBP was detected by Western blotting analysis with MBP antibody. CBS was used as a loading control (**lower panel**).

The vacuolar accumulation of free GFP, which is released from the GFP-ATG8 fusions during breakdown of the autophagic bodies measures autophagy-dependent vacuolar transport and degradation (Yoshimoto et al., 2004; Chung et al., 2010; Suttangkakul et al., 2011). To further examine whether GFP-ATG8h and GFP-ATG8i are similarly subject to autophagic degradation, transgenic seedlings of the *35S::GFP-ATG8h/35S::GFP-ATG8i* were treated in the dark in the C-deficient 1/2 MS medium to mimic the C deprivation and thus induce autophagy. As shown in Figure 9C, the C-starvation treatment significantly induced the release of the free GFP from both GFP-ATG8h and GFP-ATG8i. Treatment with ConA increased the accumulation levels of both the un-degraded GFP-ATG8h/GFP-ATG8i fusions and the released free GFP, indicating GFP-ATG8h and GFP-ATG8i are degraded in the vacuoles via autophagy pathway (Figure 9C). Similarly, transiently over-expressed CLC2-MBP via agro-infiltration in the leaves of *N. benthamiana* was also subjected to autophagic degradation under C-starvation conditions, because the release of free MBP was significantly induced and the treatment with E-64d, a Cys protease inhibitor that blocks protein degradation in the vacuole, increased the levels of both CLC2-MBP and free MBP (Figure 9D). Collectively, these data indicate that both GFP-ATG8h/GFP-ATG8i and CLC2-MBP are targeted to vacuoles for autophagic degradation and the autophagic degradation of CLC2 could be mediated by interacting with ATG8h/ATG8i (Figure 7).

## Discussion

The significance of our findings is that we provided direct genetic, biochemical and cytological evidence that partial loss of CLC function leads to the activation of SA-and H_2_O_2_-dependent cell death and immunity through compromising autophagy in Arabidopsis. CLC2 directly interacts with ATG8h and ATG8i in an AIM-and LDS-dependent manner. In addition, both CLC2 and ATG8h/ATG8i are subjected to autophagic degradation. Collectively, these results establish a direct link between the function of CLCs and the autophagy pathway.

### The functional roles of Arabidopsis CLC2 and CLC3 in SA-and H2O2-dependent cell death and its relationship with autophagy

The similarities shared between the *clc2-1clc3-1* and the *atg* mutants in the SA-and H_2_O_2_-dependent accelerated senescence, activated defense responses and enhanced resistance to biotrophic pathogens (Figure 1-4) strongly indicate a close inter-connection between the function of CLC2/CLC3 and the autophagy pathway. It has been well documented that mutations in many *ATG* genes result in over-accumulation of SA and H_2_O_2_, accelerated senescence, and excessive programmed cell death (PCD) irrespective of nutrient conditions (Doelling et al., 2002; Hanaoka et al., 2002; Xiong et al., 2007; Yoshimoto et al., 2009; Chung et al., 2010; Lai et al., 2011; Lenz et al., 2011; Wang et al., 2011). Accelerated senescence and PCD are suppressed by either degradation of SA, impairment of SA biosynthesis, or by blocking SA signaling (Yoshimoto et al., 2009; Wang et al., 2011). These results indicate that autophagy negatively regulates SA homeostasis and signaling and this negative feedback regulatory mechanism keeps senescence and immunity-related PCD under control (Yoshimoto et al., 2009). Given that loss of *CLC2* and *CLC3* compromises the autophagy pathway in Arabidopsis (Figure 5), the auto-immune phenotypes observed in the *clc2-1clc3-1* mutant (Figure 1) most likely originated from compromised autophagy.

We showed that autoimmune phenotypes of the *clc2-1clc3-1* mutant were not only rescued by loss of function of SID2/ICS1 but also by loss of function of RBOHD (Figure 2). These findings strongly indicate that both RBOHD-dependent H_2_O_2_ and SID2/ICS1-dependent SA are absolutely required for triggering the auto-immune responses in *clc2-1clc3-1* mutant plants, and the presence of either elevated SA or H_2_O_2_ alone is not sufficient to initiate and/or maintain the positive SA-ROS amplification loop (Figure 2). In support of this conclusion, it has been shown that increased ROS levels induce SA biosynthesis (Neuenschwander et al., 1995; Chamnongpol et al., 1998) and SA induces strong oxidative stress responses (Rao et al., 1999; Xue et al., 2013). The over-accumulation of ROS in the *atg2* and *atg5* mutants was significantly reduced in the *NahG* and *sid2* background plants (Yoshimoto et al., 2009). Our findings suggest that the concomitant presence of both SA and H_2_O_2_ is required for the autoimmune responses observed in the *clc2-1clc3-1* double mutant plants, and the presence of either elevated SA or H_2_O_2_ alone is not sufficient to trigger and subsequently amplify the immune response signals. The function of CLC2/CLC3 is required for attenuating the SA-ROS feed-back amplification loop (Figure 2), which might be executed through the autophagy pathway (Figure 3-5; Yoshimoto et al., 2009).

The chloroplast is the major source of SA biosynthesis in Arabidopsis (Wildermuth et al., 2001) and the PM-localized NAPDH complex is one of the primary sources of ROS induced in response to pathogen infections (Torres et al., 2002). Clathrin function is required for salt-induced RBOHD endocytosis, which is required for RBOHD-dependent ROS production (Hao et al., 2014). We reasoned that the elevated RBOHD-dependent ROS production under various stress conditions including pathogen infections (Torres et al., 2002), induces ICS1-dependent biosynthesis of SA. As a result, both elevated primary ROS and SA induce secondary ROS production from the organelles such as chloroplasts, mitochondria, and peroxisomes (Mittler, 2017), which subsequently triggers the positive SA-ROS amplification loop. Damaged or dysfunctional organelles under severe stress conditions are eliminated by the selective autophagy processes such as chlorophagy (Ishida et al., 2008; Izumi et al., 2010; Xie et al., 2015), mitophagy (Li et al., 2014), and pexophagy (Young and Bartel, 2016; Luo et al., 2018), which keep the positive SA-ROS amplification loop in check. In agreement, we showed that the ROS level was significantly higher in the chloroplasts of *clc2-1clc3-1* and *atg2-1* than in that of Col-0 (Figure 6), indicating that the enhanced accumulation of both SA and H_2_O_2_ in the *clc2-1clc3-1* and the *atg2-1* mutant could be due to over-accumulation of dysfunctional organelles as a result of impaired autophagy pathway. Autophagy may play a critical role in dampening the positive ROS-SA amplification loop by reducing secondary SA and ROS production through eliminating damaged chloroplasts, peroxisomes, or mitochondria. The fact that the *clathrin heavy chain 2* (*CHC2*) mutants displayed a similar autoimmune phenotype (Wu et al., 2015) implies that loss of CHC2 may similarly result in activated immune responses through impairing the autophagy pathway. It seems that a closely inter-connected and mutually regulated relationship between CME and autophagy exists in Arabidopsis.

### The potential roles of clathrin in autophagy

The number of autophagosomes and/or autophagic bodies was significantly reduced in the *clc2-1clc3-1* mutant in the presence of E-64d, a Cys protease inhibitor that blocks protein degradation in the vacuole (Figure 5A, 5B). This could be due to impaired formation of autophagosomes, reduced delivery of autophagosomes into vacuoles, compromised degradation of autophagic bodies inside vacuole, or any combination of these possibilities (Zhuang et al., 2015). The fact that the C starvation-induced accumulation levels of both ATG8 and NBR1 in the leaves of *clc2-1clc3-1* mutant were not increased to the extent observed in that of *atg2-2* in the presence of E-64d (Fig. 5C) suggest that, in addition to the degradation in the vacuoles, the biogenesis of autophagosomes is also compromised in *clc2-1clc3-1*. The functional connection of clathrin with autophagy has previously been reported in mammals. In human cells, autophagy contributed to connexin 31.1 (Cx31.1) degradation, and clathrin might be involved in the autophagy of Cx31.1 (Zhu et al., 2015). Clathrin, AP2, AP4, and PtdIns(4,5)P2 directly mediate budding of the autolysosome in rat (Rong et al., 2012). The human clathrin adaptor protein AP1 and clathrin localized to starvation-and rapamycin-induced autophagosomes and dysfunction of the AP1-dependent clathrin coating at the TGN prevented autophagosome formation (Guo et al., 2012). In mammalian cells, CHC and AP-2 interact with Atg16L1 and depletion of CHC and AP-2 moderately, but significantly, reduces autophagy initiation and the formation of pre-autophagosome structures (Ravikumar et al., 2010; Lamb et al., 2012).

Although indirect evidence implied that clathrin may play a role in autophagy in plants, direct evidence showing that clathrin plays a role in autophagy is still lacking. It has been shown recently that a SH3-domain containing protein 2 (SH3P2), a BAR (Bin-Amphiphysin-Rvs) domain protein, participates in autophagosome formation in Arabidopsis through interacting with ATG8e/8f, PI3P and associating with PI3K (Zhuang et al., 2012; 2013; Gao et al., 2015). SH3P2 localizes to the PM, associates with CCVs, co-localizes with CLC-labeled structures and coimmunoprecipitates with CHC (Nagel et al., 2017). Based on these observations, clathrin is proposed to play a role in the initiation of autophagosome formation, membrane expansion or maturation or cargo sorting in Arabidopsis (Zhuang et al., 2012; 2013; 2015; Gao et al., 2015). Arabidopsis AtEH1/Pan1 and AtEH2/Pan1 proteins, the subunits of endocytic TPLATE complex (TPC), are involved in actin cytoskeleton regulated autophagy (Wang et al., 2019). When autophagy is induced, AtEH/Pan1 proteins recruit TPC and AP-2 subunits, CHC, and ARP2/3 proteins for the formation of the autophagosomes through interaction with F-actin and VAP27-1 at ER (Endoplasmic reticulum)-PM contact sites (Wang et al., 2019). However, comparison of the mutants in the CME pathway to *atg* mutants was not included in this study. Therefore, it is unclear whether these mutants share similar morphological phenotype and biochemical characteristics with *atg* mutants. In addition, direct genetic evidence for the involvement of CLCs or CHCs in the SH3P2-or AtEH/Pan1-mediated autophagy was not provided (Zhuang et al., 2012; 2013; 2015; Gao et al., 2015; Wang et al., 2019).

### The functional relevance of the CLC2-ATG8h/ATG8i interactions

The unique feature for the ATG8h-i group is that they have an exposed catalytic glycine residue at the C terminus (Figure 5S), and do not require cleavage by the ATG4 protease for activation (Seo et al., 2016; Kellner et al., 2017). The cargoes targeted by selective autophagy are usually recognized by cargo receptors that interact with membrane-anchored ATG8s through the ATG8-interacting motifs (AIM) present on the cargo receptors (Noda et al., 2010; Birgisdottir et al., 2013). The ATG8h has been reported to interact with ER/plastid bodies-localized ATG8-interacting proteins 1 (ATI1) (Avin-Wittenberg et al., 2012). These interactions function to target the damaged ER/plastid and/or ER/plastid proteins to the vacuoles for degradation (Michaeli et al., 2014). Therefore, it is plausible that the CLC2-ATG8h/ATG8i–ATI1/2 interactions redirect the ER/plastid bodies to the sites of autophagosome formation for the membrane components. The tomato ATG8h interacts with AvrPto-dependent Pto-interacting protein 3 (Adi3), an AGC protein kinase that function to suppress programmed cell death (PCD) (Devarenne, 2011). In another case, the interaction of the nucleus-localized C1, a replication initiator protein of Tomato leaf curl Yuannan virus (TLCYnY), with *Nb*ATG8h leads to the translocation of the C1 protein from the nucleus to the cytoplasm and results in its degradation by selective autophagy in an AIM-dependent manner (Li et al., 2019). These results indicate that ATG8h participates in different biological processes through interacting with different proteins.

It has been shown that TPLATE and TML, two subunits of the TPC at the PM (Gadeyne et al., 2014), interacts with CLC2 (Van Damme et al., 2011; Gadeyne et al., 2014). Here, we showed that CLC2 interacts with ATG8h/ATG8i in an AIM1-and LDS-dependent manner (Figure 7 and Figure 8). The functional relevance of the CLC2-ATG8h/ATG8i interaction was reflected by the facts that both GFP-ATG8h/ATG8i and CLC2 were subjected to autophagic degradation in the vacuoles (Figure 9). These results suggest that CLC2 may serve as an autophagic receptor to selectively mediate the autophagic degradation of CLC2 itself and CCVs including EEs/TGNs, which are coated with clathrins (Viotti et al., 2010; Ito et al., 2012; Konopka et al., 2008) (Figure 10, Pathway 1), which in turn may regulate the progression of autophagy (Zhuang and Jiang, 2014; Zhuang et al., 2015). However, the CLC2-ATG8h/ATG8i interactions may also provide a means to deliver the CCVs carrying lipids/phospholipids (Lipid nanodrops), PM proteins or other necessary components to the sites of autophagosome biogenesis (Figure 10, Pathway 2; Zhuang et al., 2013; 2015; 2018; Gao et al., 2015; Gomez et al., 2022). The significantly reduced number of autophagic bodies and autophagic flux observed in the root cells of *clc2-1clc3-1* mutant seedlings under C starvation conditions (Figure 5A and 5B) supports this statement. Given that clathrin is also involved in post-Golgi trafficking (Reynolds et al., 2018; Yan et al., 2021), the CLC2-ATG8h/ATG8i interactions may provide membrane components required for the biogenesis of autophagosome from the ER through secretory pathway (Ravikumar et al., 2010; Zhuang et al., 2013; 2015; 2018).

**Figure 10.**
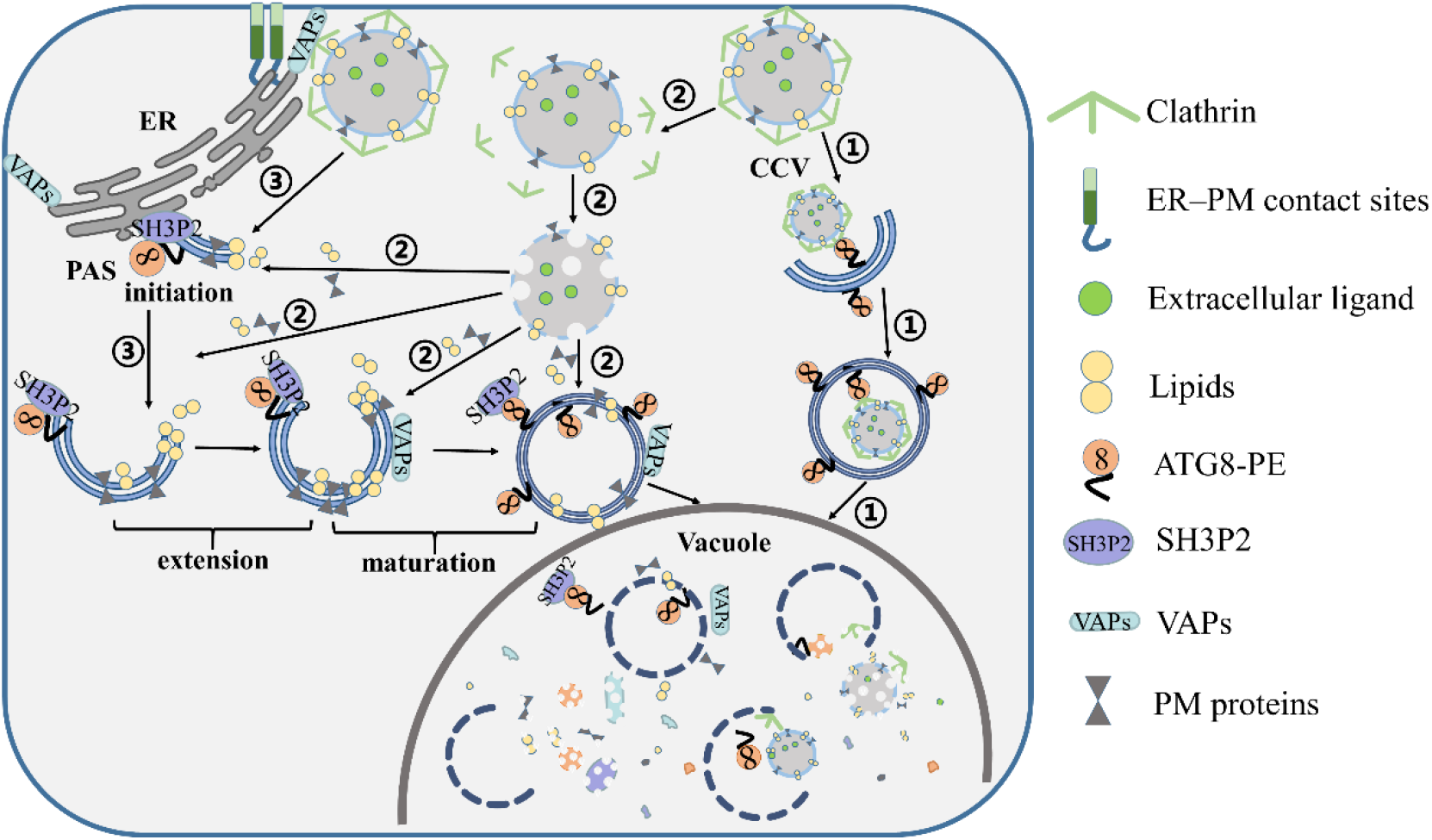
A proposed working model for CLC2 or CME in participating autophagy pathway. Pathway 1: CLC2 serves as a cargo receptor to mediate the selective degradation of CLC2, CCV or TGN/EE through directly interacting with ATG8h/ATG8i; Pathway 2: CCVs carry membrane proteins, various lipids such as PIPs and other components required for the biogenesis of autophagosomes to promote initiation, extension, expansion and maturation of autophagosomes. Pathway 3: Clathrin indirectly participates in the biogenesis of autophagosomes through interacting with the key proteins, such as SH3P2 and VAMP-associated-proteins (VAP), in the autophagy pathway.

In addition to directly participating in autophagy via interacting with ATG8h/ATG8i, CLC2 may also be involved in autophagy indirectly. The ER membrane-associated VAMP-associated-proteins (VAP) establish a close association with endosomes and contact the PM at EPCSs (Stefano et al., 2018). Arabidopsis VAP27-1 and VAP27-3 interact with clathrin and bind phosphatidylinositolphosphate lipids (PIPs) including PI3P (Stefano et al., 2018), which also decorates autophagosome membranes (Li and Viestra, 2012), raising a possibility that VAP27 proteins bridge the ER with autophagosome membranes likely through their interaction with CLCs or CHCs (Figure 10, Pathway 3; Stefano et al., 2018). ATG8h/8i-CLC2-VAP27s-AtEH/Pan1 four-way interactions (Stefano et al., 2018; and Wang et al., 2019) could be involved in the AtEHs/Pan1-mediated autophagosome formation at EPCS (Wang et al., 2019).

SH3P2 interacts with CLC and CHC (Adamowski et al., 2018; Nagel et al., 2017), respectively, and translocates to the phagophore assembly site/preautophagosome structure (PAS) upon autophagy induction (Zhuang et al. 2015). In addition, SH3P2-GFP decorates ER-derived omegasome-like structures (Zhuang et al. 2015). Furthermore, SH3P2 also interacts with all ATG8 isoforms in Arabidopsis and an AIM-like motif within the SH3 domain, which is essential for the SH3P2-ATG8f interaction (Zhuang et al, 2013; Sun et al., 2022). Therefore, clathrin can be associated with autophagosomes through interacting with ATG8h/ATG8i directly (Fig. 7) or through interacting with other ATG8s indirectly (Zhuang et al., 2013; Sun et al., 2022). The CLC2-ATG8h/ATG8i interaction may deliver the PM-derived lipids and other required component via TGN/EE to the initiating or expanding autophagosomes (Figure 10, Pathway 2), whereas the CLC-SH3P2-ATG8a-8i interactions may facilitate delivering of both PM-and ER/EPCS-derived membrane or protein components to the sites of autophagosome biogenesis (Figure 10, Pathway 3) Nonetheless, our data indicate that CLC2 participates in the autophagy pathway through directly interacting with ATG8h/8i, establishing a previously unidentified functional link between CME and autophagy, two major conserved intracellular degradation pathways across kingdoms.

## Materials and Methods

The following T-DNA mutants were either from the ABRC or the Nottingham Arabidopsis Stock Centre: *atg2-1* (SALK_076727) (Yoshimoto et al., 2009), *atg2-2* (Wang et al., 2011), *atg5* (SALK_020601), *atg7* (SAIL_11_H07). Homozygous T-DNA insertion mutants were identified by genomic PCR (Alonso et al, 2003). The transgenic Arabidopsis plants expressing GFP-ATG8e in the background of Col-0 and *atg2-2* were described (Wang et al., 2011).

### Generation of double or triple mutants

*clc2-1clc3-1* (Wang et al., 2013) mutant was crossed with *sid2-2* (Wildermuth et al., 2001) and *rbohD* (Torres et al., 2002), *atg2-1*, *atg5* and *atg7* to generate respective triple mutants. The primers used for identifying the triple mutants by genomic PCR are CLC2-CS808198/SALK_016049-F; CLC2-CS808198/SALK_016049-R; CLC2-GK-T-DNA-LB1; CLC3-CS100219-F; CLC3-CS100219-R; CLC3-sgtDs3’-1;rbohD-dSpm1; rbohD-LP; rbohD-RP; sid2XS691-LP; sid2XS692-RP and sid2-LBal. The primer sequence information listed in Supplemental Table 1.

### SA Quantification

SA was quantified using an Agilent 1260 HPLC system (Agilent Technologies) with a diode array detector and a fluorescence detector and a column as described previously (Zhang et al., 2017).

### H_2_O_2_ quantification by DAB Staining

H_2_O_2_ was visualized by a DAB staining procedure (Sigma-Aldrich). Detached leaves were placed in DAB solution (1 mg mL−1 DAB, pH 5.5) for 2 h followed by clearance in 96% (v/v) boiling ethanol for 10 min. H_2_O_2_ production was detected as a reddish-brown precipitate on the cleared leaves (Ren et al., 2002).

### Trypan blue staining of dead cells

Plant cell death and fungal growth were visualized using trypan blue staining as described (Wang et al., 2011).

### Chlorophyll content measurements

Chlorophyll contents were measured as described previously (Qi et al., 2017). Briefly, Arabidopsis leaves were extracted by incubation in 1 mL of N,N-dimethylformamide for 48-h in the dark at 4°C. Absorbance at 664-and 647-nm was read, respectively, and the total chlorophyll content was calculated and normalized to fresh weight (gram) per sample.

### Powdery mildew infections

*G. cichoracearum* strain UCSC1 was maintained by inoculating on *sid2* mutant plants. 6-week-old plants were inoculated with *G. cichoracearum* strain UCSC1 and the symptoms were scored at 7 days-post-inoculation (dpi) as described (Wang et al., 2011). Trypan blue assay was performed to monitor the fungal structures and dead cells. The number of conidiophores per colony was counted at 7 dpi. At least 20 colonies were counted for each genotype in one experiment. The experiments were repeated three times with similar results.

### Pseudomonas syringae pv. tomato DC3000 (Pst DC3000) infection

The *Pst* DC3000 infection and HR tests were performed as described (Heese et al., 2007). The bacterial culture of OD_600_=1 was diluted to 10^-4^ in 10 mM MgCl2. Four-week-old plants grown under short day condition (8-h light/16-h dark) were infiltrated with the bacterial suspensions and bacterial growth was monitored at the dpi as indicated.

### RNA extraction and RT-qPCR

RNA isolation and RT-qPCR were performed as described elsewhere (Xu et al., 2018). The RT-qPCRs were performed using an ABI550 Real-Time PCR machine (Applied Biosystems, Thermo Fisher Scientific) and the 2x SYBR Green qPCR Mix (Aidlab, Beijing, China). The primers used for RT-qPCR are *PR1*-F; *PR1*-R; *ACTIN2*-F; *ACTIN2*-R. The sequence information of the primers is listed in Supplemental Table 1.

### Western blotting analysis

Proteins were prepared from leaves collected from 4-week-old Col-0, *clc2-1clc3-1* and *atg2-1* plants treated in the dark for different time and the immunoblots were performed as previously described (Qi et al., 2017). Anti-NBR1 (Agresera, AS142805), anti-ATG8 (Agresera, AS142769), anti-ATG5 (Agresera, AS153060), anti-GFP (Santa Cruz, SC-81045) and anti-MBP (CWBIO, 01254) were used for Western blotting.

### Yeast Two-Hybrid Assays

The full length coding sequences of the *CLC2*, *CLC3* or *ATG8a*-*ATG8i* were amplified from the cDNA transcribed from the mRNA extracted from Arabidopsis seedlings and subsequently cloned into pGBKT7 (BD) or pGADT7 (AD) vector (Clontech, Mountain View, CA), respectively. The primers used for the amplifications listed in Supplemental Table 1. Positive interactions were confirmed by growth of the co-transformed yeast on selective medium lacking Trp, Leu, His and Ade.

### Site-directed Mutagenesis

The full length cDNA of *CLC2* was firstly cloned into a pMD^TM^20 T-vector (Takara Bio, San Jose, CA). The *CLC2* containing AIM1 deleteion and AIM2 mutation as well as LDS mutated ATG8h were generated respectively by a Mut Express^®^ II Fast Mutagenesis Kit V2 (Vazyme, Nanjing, China) using primers containing the intended mutations. The primers used for the amplifications listed in Supplemental Table 1.

### Luciferase Complementation Imaging Assay

The CLC2, CLC2-AIM1, CLC2-AIM2, CLC2-AIM3 and ATG8h or ATG8i full length coding sequences were cloned into Firefly Luciferase Complementation Imaging vectors 35S::nLuc and 35S::cLuc (Chen et al., 2008), respectively, by digestion with BamHI + SalI or KpnI to generate constructs fused either with N-terminal or C-terminal half of luciferase. The primers used for the amplifications listed in Supplemental Table 1. The generated CLC2 and ATG8h fusion construct pairs were co-injected into *N. benthamiana* leaves, respectively, via agro-infiltration. At 2 days post infiltration, the abaxial surfaces of the infiltrated leaves was sprayed with 1 mmol/L D-luciferin (MedChemExpress, Shanghai, China) and kept in dark for 7 min to quench the fluorescence followed by capturing the LUC image with a low-light cooled CCD imaging apparatus (Tanon 6600, Biotanon, Shanghai, China).

### Confocal microscopy

The transgenic Col-0 and *atg2-2* plants expressing GFP-ATG8e was generated previously (Wang et al., 2011). The *clc2-1clc3-1* line expressing GFP-ATG8e (*clc2-1clc3-1*/GFP-ATG8e) was created by crossing with Col-0/GFP-ATG8e and screened in T2 and T3 populations by genomic PCR using following primers: CLC2-CS808198/SALK_016049-F; CLC2-CS808198/SALK_016049-R; CLC2-GK-T-DNA-LB1; CLC3-CS100219-F; CLC3-CS100219-R; CLC3-sgtDs3’-1; GFP-F; and ATG8e-R. The sequence information of the primers is listed in Supplemental Table 1.

Five-day-old transgenic seedlings grown on the 1/2 MS medium with or without sucrose were treated under dark for 2 days followed by incubation with or without E-64d, a Cys protease inhibitor that blocks protein degradation in the vacuole (Yoshimoto et al., 2004). The visualization of autophagic bodies represented by the expression of the GFP-ATG8e fusion protein in the primary roots of was performed using a confocal microscopy (Zeiss, LSM880, Oberkochen, Germany) with 488 nm laser light excitation and a 490–540 nm band-pass filter set as described (Wang et al., 2011). Image processing was performed according to the manufacturer’s manual.

CLC2-mKO, GFP-ATG8h and GFP-ATG8i fusions were amplified by PCRs or over-lapping PCRs using the primers listed in the Supplemental Table 1. The 35S::CLC2-mKO, 35S::GFP-ATG8h and 35S::GFP-ATG8i constructs were made by cloning the amplified PCR products into pE1776 vector (Leister et al., et al., 2005). The CLC2-mKO+GFP-ATG8h or CLC2-mKO+GFP-ATG8i fusion constructs were infiltrated into *N. benthamiana* leaves as described (Liu et al., 2005). Images were captured with a confocal laser-scanning microscope (Carl Zeiss, LSM 880).

## Author contributions

J.Z.L., S.A.W. and J.P. conceived the study and designed the experiments. H.J.L., J.R., L.Z., N.N.W., W.X.W., M.N., N.C. conducted the experiments. J.Z.L. analyzed the data. J.Z.L. wrote the manuscript and S.A.W., P.A.N. and N.H.C. revised the manuscript.

## Acknowledgments

We dedicate this paper in memory of our friend and colleague, Prof. Jianwei Pan. His passion for science and fighting spirit will inspire us for years to come. We thank Prof. Dingzhong Tang for kindly providing *atg2-1* mutant, GFP-ATG8e/Col-0, GFP-ATG8e/*atg2-2* transgenic lines and *G. cichoracearum* strain UCSC. We also thank Profs. Brian J. Staskawicz, Sebastian Y. Bednarek, Kewei Zhang and Chao Wang for generously sharing materials, constructs and vectors.

