## Supplemental Figures for "Clathrin Light Chains are essential in negative regulation of cell death and immunity in Arabidopsis through interacting with autophagy pathway"

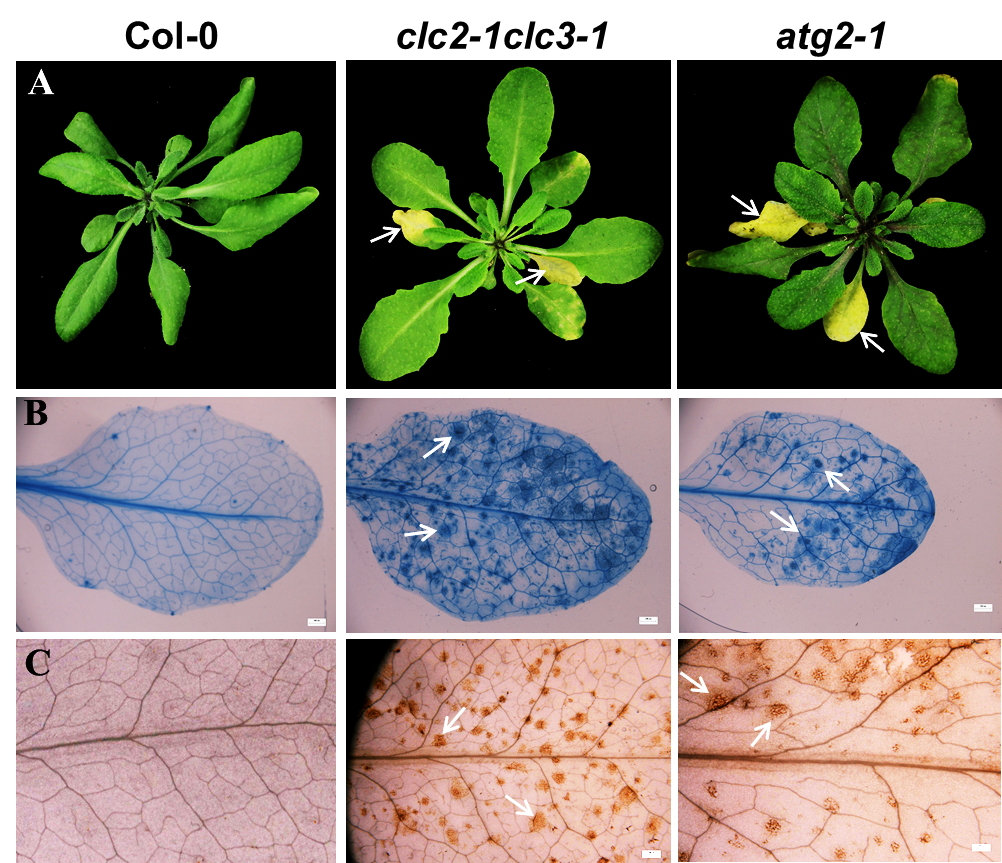

**Figure 1S.** **The** ***clc2-1clc3-1* double mutant and the *atg2-1* mutant share similar auto-immune phenotypes.** (**A**) The *clc2-1clc3-1* double mutant and the *atg2-1* mutant displayed similar accelerated senescence and chlorotic cell death at 4 weeks after sowing. (**B**) Cell death was visualized by trypan blue staining on the leaves of *clc2-1clc3-1* double mutant and *atg2-1* mutant plants (see arrows) but not on the leaves of the Col-0 plants. (**C**) Enhanced accumulation of H_2_O_2_ visualized by DAB staining was observed on the leaves of the *clc2-1clc3-1* double mutant and the *atg2-1* mutant plants (see arrows) but not on the leaves of Col-0 plants.

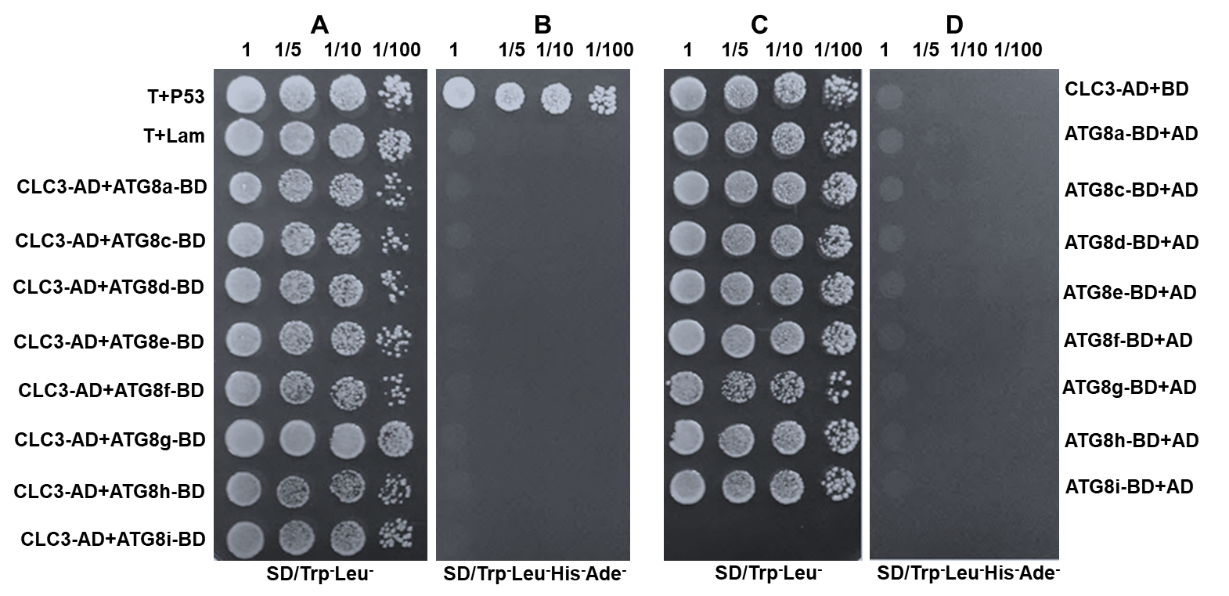

**Figure 2S**. **CLC3 does not interact with ATG8s in a yeast two-hybrid assay.** Yeast cells in **A** and **C** were grown on SD/Trp^-^Leu^-^ medium; Yeast cells in **B** and **D** were selected on SD/Trp^-^Leu^-^His^-^Ade^-^ medium. CLC3 interacts with none of the ATG8s tested. CLC3-AD also does not interact with empty BD vector. T-antigen and p53 were included as a positive interaction control, and Lam and T-antigen were included as a negative interaction control; AD: activation domain; BD: binding domain. The numbers on top of the images are the diluting factors.

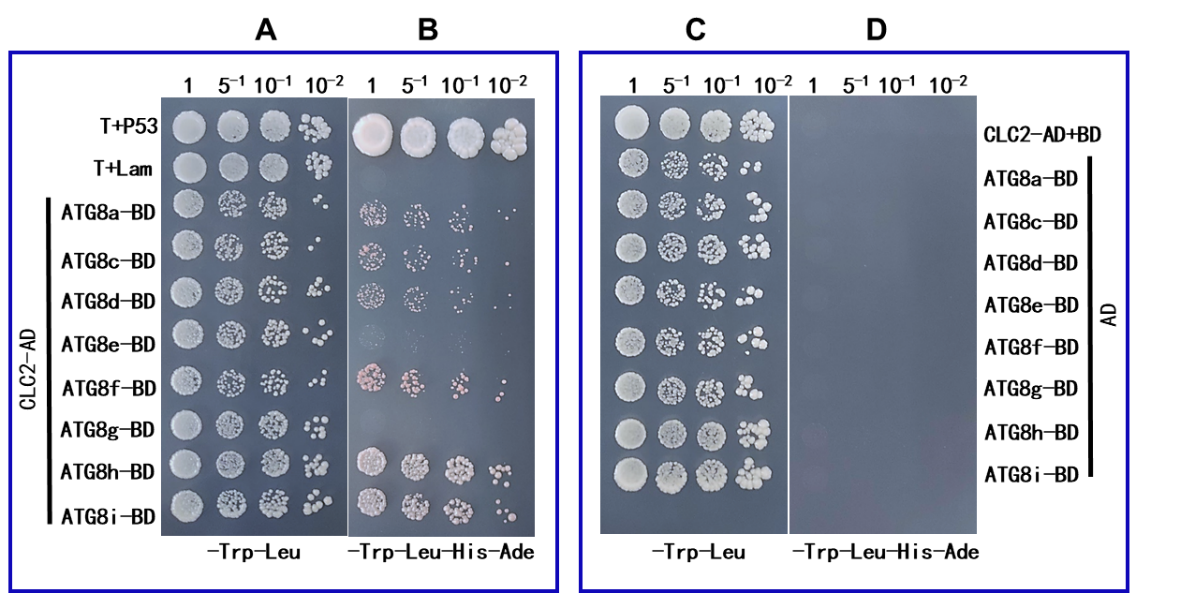

**Figure 3S. CLC2 strongly interacts with ATG8h and ATG8i and only weakly with ATG8a, ATG8c, ATG8d and ATG8f in the yeast two-hybrid assay.** Yeast cells in **A** and **C** were grown on SD/Trp^-^Leu^-^ medium; Yeast cells in **B** and **D** were selected on SD/Trp^-^Leu^-^His^-^Ade^-^ medium. The images were taken at 7 days post inoculation. T-antigen and p53 were included as a positive interaction control, and Lam and T-antigen were included as a negative interaction control; AD: activation domain; BD: binding domain. The numbers on top of the images are the diluting factors.

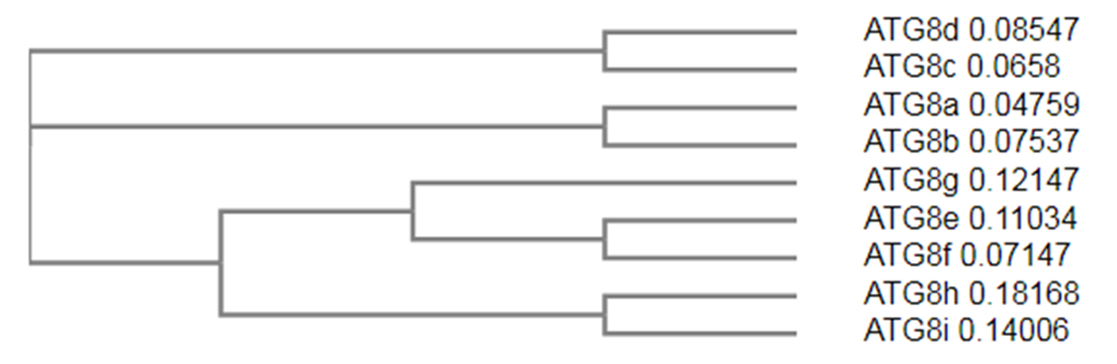

**Figure 4S. The phylogenetic tree of the Arabidopsis ATG8 homologs.** The tree was generated by using ClustalW. ATG8h and ATG8i are clustered in the same clades.

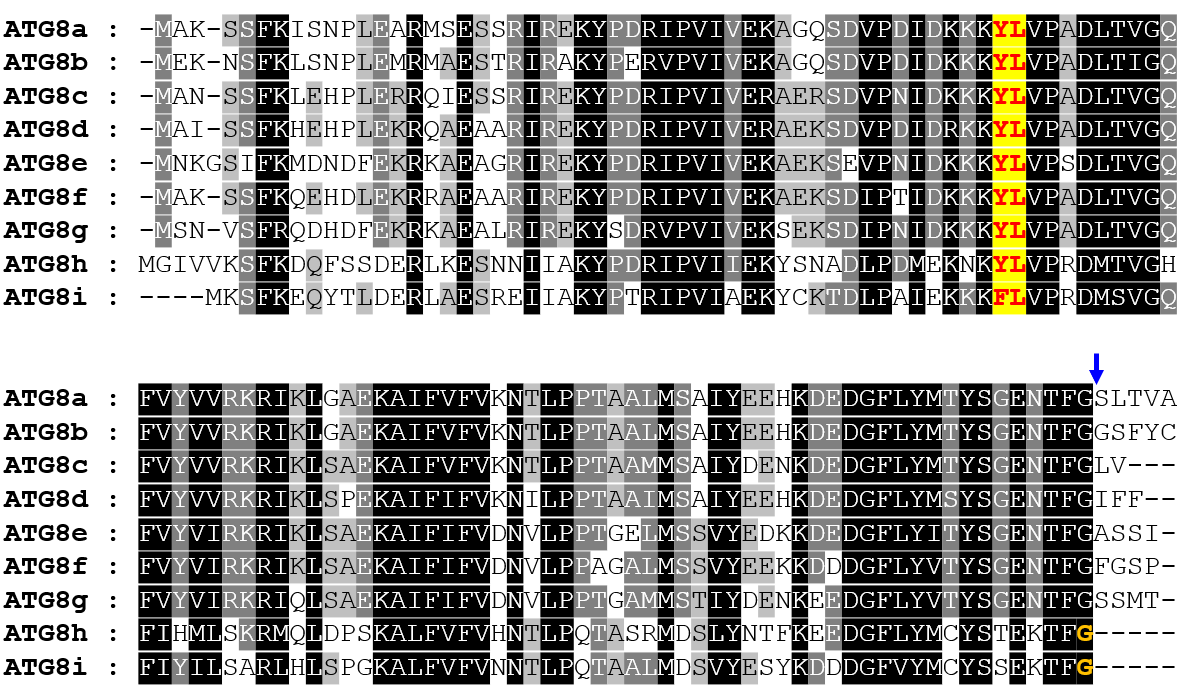

**Figure 5S. The amino acid sequence alignment of ATG8 homologs**. The alignment was performed using MEGA11. The glycine residue of ATG8a-ATG8g, the cleavage site of ATG4 protease for activation of ATG8 homologs, is marked by blue arrow. The exposed catalytic glycine residue at the C termini of ATG8h and ATG8i is shown in orange. The conserved LIR/AIM docking site (LDS) presented in the ATG8 protein family is highlighted in yellow.

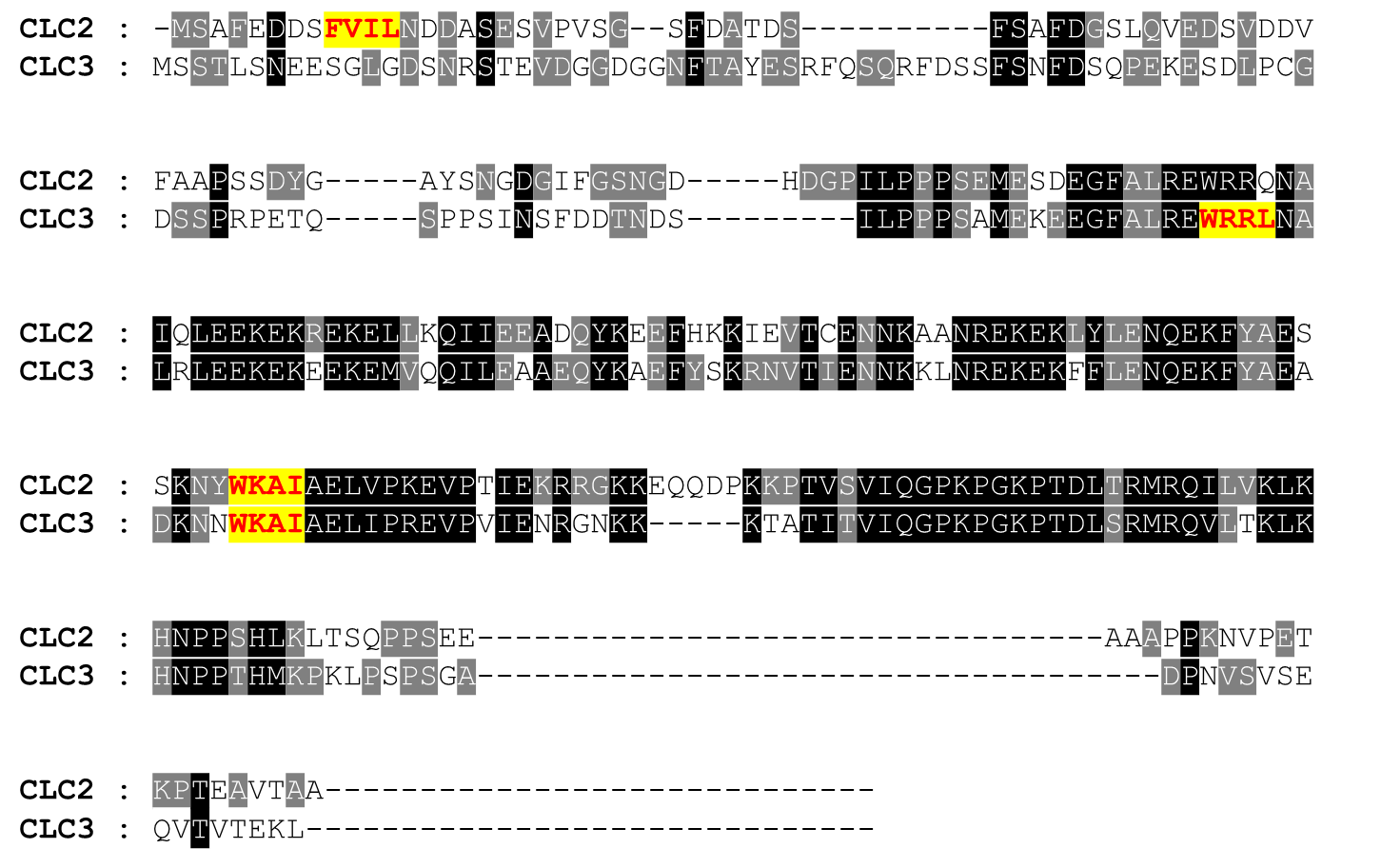

**Figure 6S. The amino acid sequence alignment of CLC2 (At2g40060) and CLC3 (At3g51890)**. The alignment was performed using MEGA11. The putative ATG8-interacting motifs (AIMs) are highlighted in yellow. The AIMs are predicted by using High-Fidelity AIM (hfAIM) (<http://bioinformatics.psb.ugent.be/hfAIM/>) (Xie et al., 2016, Autophagy 12: 876-887).
