## Supplemental Table 1 for "Clathrin Light Chains are essential in negative regulation of cell death and immunity in Arabidopsis through interacting with autophagy pathway"

**Supplemental Table 1. The primers used in this study**

| **Constructs** | **Accession #** | **Primer names** | **Primer sequences** |
| --- | --- | --- | --- |
| CLC2-AD | AT2G40060.1 | CLC2-NdeI-F | 5'-3'aaa**CATATGA**TGTCTGCCTTTGAAGACG |
|  |  | CLC2-ECORI-R | 5'-3'ttt**GAATTC**TTAAGCAGCAGTAACTGCCT |
| CLC2-BD | AT2G40060.1 | CLC2-NdeI-F | 5'-3'aaa**CATATGA**TGTCTGCCTTTGAAGACG |
|  |  | CLC2-EcoRI-R | 5'-3'ttt**GAATTC**TTAAGCAGCAGTAACTGCCT |
| CLC3-AD | AT3G51890.1 | CLC3-EcoRI-F | 5'-3'aaa**GAATTC**ATGTCGTCAACCTTGAGCA |
|  |  | CLC3-BamHI-R | 5'-3'ttt**GGATCC**CTACAACTTCTCTGTAACTGTGACC |
| CLC3-BD | AT3G51890.1 | CLC3-EcoRI-F | 5'-3'aaa**GAATTC**ATGTCGTCAACCTTGAGCA |
|  |  | CLC3-BamHI-R | 5'-3'ttt**GGATCC**CTACAACTTCTCTGTAACTGTGACC |
| ATG8a-AD | AT4G21980.1 | ATG8a-NdeI-F | 5'-3'aaa**CATATG**ATGGCTAAGAGTTCCTTCAAGA |
|  |  | ATG8a-BamHI-R | 5'-3'ttt**GGATCC**TCAAGCAACGGTAAGAGATCC |
| ATG8a-BD | AT4G21980.1 | ATG8a-NdeI-F | 5'-3'aaa**CATATG**ATGGCTAAGAGTTCCTTCAAGA |
|  |  | ATG8a-BamHI-R | 5'-3'ttt**GGATCC**TCAAGCAACGGTAAGAGATCC |
| ATG8c-AD | AT1G62040.1 | ATG8c-NdeI-F | 5'-3'cgc**CATATG**ATGGCTAATAGCTCTTTCAAGTTG |
|  |  | ATG8c-BamHI-R | 5'-3'ttt**GGATCC**TTAAACCAAACCAAAGGTGTTCT |
| ATG8c-BD | AT1G62040.1 | ATG8c-NdeI-F | 5'-3'cgc**CATATG**ATGGCTAATAGCTCTTTCAAGTTG |
|  |  | ATG8c-BamHI-R | 5'-3'ttt**GGATCC**TTAAACCAAACCAAAGGTGTTCT |
| ATG8d-AD | AT2G05630.1 | ATG8d-NdeI-F | 5'-3'aaa**CATATG**ATGGCGATTAGCTCCTTCAA |
|  |  | ATG8d-BamHI-R | 5'-3'ttt**GGATCC**TTAGAAGAAGATCCCGAACGTG |
| ATG8d-BD | AT2G05630.1 | ATG8d-NdeI-F | 5'-3'aaa**CATATG**ATGGCGATTAGCTCCTTCAA |
|  |  | ATG8d-BamHI-R | 5'-3'ttt**GGATCC**TTAGAAGAAGATCCCGAACGTG |
| ATG8e-AD | AT2G45170.1 | ATG8e-NdeI-F | 5'-3'aaa**CATATG**ATGAATAAAGGAAGCATCTTTAAGATG |
|  |  | ATG8e-BamHI-R | 5'-3'ttt**GGATCC**TTAGATTGAAGAAGCACCGAATG |
| ATG8e-BD | AT2G45170.1 | ATG8e-NdeI-F | 5'-3'aaa**CATATG**ATGAATAAAGGAAGCATCTTTAAGATG |
|  |  | ATG8e-BamHI-R | 5'-3'ttt**GGATCC**TTAGATTGAAGAAGCACCGAATG |
| ATG8f-AD | AT4G16520.1 | ATG8f-NdeI-F | 5'-3'aaa**CATATG**ATGGCAAAAAGCTCGTTCAA |
|  |  | ATG8f-BamHI-R | 5'-3'ttt**GGATCC**TTATGGAGATCCAAATCCAAATGT |
| ATG8f-BD | AT4G16520.1 | ATG8f-NdeI-F | 5'-3'aaa**CATATG**ATGGCAAAAAGCTCGTTCAA |
|  |  | ATG8f-BamHI-R | 5'-3'ttt**GGATCC**TTATGGAGATCCAAATCCAAATGT |
| ATG8g-AD | AT3G60640.1 | ATG8g-NdeI-F | 5'-3'aaa**CATATG**ATGAGTAACGTCAGCTTCAGG |
|  |  | ATG8g-BamHI-R | 5'-3'ttt**GGATCC**TTAAGTCATTGACGATCCAAAAGT |
| ATG8g-BD | AT3G60640.1 | ATG8g-NdeI-F | 5'-3'aaa**CATATG**ATGAGTAACGTCAGCTTCAGG |
|  |  | ATG8g-BamHI-R | 5'-3'ttt**GGATCC**TTAAGTCATTGACGATCCAAAAGT |
| ATG8h-AD | AT3G06420.1 | ATG8h-EcoRI-F | 5'-3'aaa**GAATTC**ATGGGGATTGTTGTCAAGTCT |
|  |  | ATG8h-BamHI-R | 5'-3'ttt**GGATCC**TTAGCCGAAAGTTTTCTCGGT |
| ATG8h-BD | AT3G06420.1 | ATG8h-EcoRI-F | 5'-3'aaa**GAATTC**ATGGGGATTGTTGTCAAGTCT |
|  |  | ATG8h-BamHI-R | 5'-3'ttt**GGATCC**TTAGCCGAAAGTTTTCTCGGT |
| ATG8i-AD | AT3G15580.1 | ATG8i-NdeI-F | 5'-3'aaa**CATATG**ATGAAATCGTTCAAGGAACAATAC |
|  |  | ATG8i-BamHI-R | 5'-3'ttt**GGATCC**TCAACCAAAGGTTTTCTCACTG |
| ATG8i-BD | AT3G15580.1 | ATG8i-NdeI-F | 5'-3'aaa**CATATG**ATGAAATCGTTCAAGGAACAATAC |
|  |  | ATG8i-BamHI-R | 5'-3'ttt**GGATCC**TCAACCAAAGGTTTTCTCACTG |
| CLC2-AIM1 | AT2G40060.1 | CLC2-AIM1-Mu1-F | TCCGCCGTCATAGCCAACGATGATGCGTCTGAGTCT |
|  |  | CLC2-AIM1-Mu1-R | TTGGCTATGACGGCGGAATCGTCTTCAAAGGCAGAC |
| CLC2-AIM2 | AT2G40060.1 | CLC2-AIM2-Mu2-F | GAAGCGAGAAGAGCAAATGCAATTCAACTTGAGGAGAA |
|  |  | CLC2-AIM2-Mu2-R | TTTGCTCTTCTCGCTTCTCTAAGAGCAAATCCCTCATCT |
| CLC2-AIM3 | AT2G40060.1 | CLC2-AIM3-Mu3-F | ATTACGCGAAGGCAGCAGCAGAGCTAGTTCCTAAAGAAGTTCC |
|  |  | CLC2-AIM3-Mu3-R | TGCTGCCTTCGCGTAATTCTTGCTGGATTCCGCG |
| cLuc-CLC2 | AT2G40060.1 | cLuc-CLC2-Kpn1-F | AAAGGTACCATGTCTGCCTTTGAAGACGAT |
|  |  | cLuc-CLC2-Kpn1-R | TTTGGTACCTTAAGCAGCAGTAACTGCCT |
| ATG8h-nLuc | AT3G06420.1 | ATG8h-nLuc-BamHI-F | AAAGGATCCATGGGGATTGTTGTCAAGTCTTTCAAG |
|  |  | ATG8h-nLuc-Sal1-R | TTTGTCGACACCAAAGGTTTTCTCACTGCTA |
| ATG8i-nLuc | AT3G15580.1 | ATG8i-nLuc-BamHI-F | AAAGGATCCATGAAATCGTTCAAGGAACAATACACG |
|  |  | ATG8i-nLuc-Sal1-R | TTTGTCGACACCAAAGGTTTTCTCACTGCTA |
| cLuc-CLC2-AIM1/2/3 | AT2G40060.1 | cLuc-CLC2-KpnI1-F | AAAGGTACCATGTCTGCCTTTGAAGACGAT |
|  |  | cLuc-CLC-KpnI1-R | TTTGGTACCTTAAGCAGCAGTAACTGCCT |
| CLC2-AIM1/2/3-AD | AT2G40060.1 | CLC2-NdeI-F | 5'-3'aaa**CATATGA**TGTCTGCCTTTGAAGACG |
|  |  | CLC2-ECORI-R | 5'-3'ttt**GAATTC**TTAAGCAGCAGTAACTGCCT |
| ATG8h-LDS-Mu | AT3G06420.1 | ATG8h-LDS-Mu-F | AAGAACAAAGCCGCGGTCCCACGAGACATGACTGTTG |
|  |  | ATG8h-LDS-Mu-R | GACCGCGGCTTTGTTCTTCTCCATGTCTGGCAG |
| ATG8h-LDS-Mu-BD | AT3G06420.1 | ATG8h-EcoRI-F | 5'-3'aaa**GAATTC**ATGGGGATTGTTGTCAAGTCT |
|  |  | ATG8h-BamHI-R | 5'-3'ttt**GGATCC**TTAGCCGAAAGTTTTCTCGGT |
| ATG8h-LDS-Mu-nLuc | AT3G06420.1 | ATG8h-nLuc-BamHI-F | TTTGGATCCATGGGGATTGTTGTCAAGTCTTTCAAG |
|  |  | ATG8h-nLuc-Sal1-R | TTTGTCGACACCAAAGGTTTTCTCACTGCTA |
| CLC2-mKO-1776 |  | CLC2-mKO-Kpn1-F | 5'-3'aaaGGTACCATGTCTGCCTTTGAAGACGAT |
|  |  | mKO-CLC2-Sma1-R | 5'-3'tttCCCGGGTCAGGAATGAGCTACTGCATC |
| GFP-ATG8e-1776 |  | GFP-ATG8e-Kpn1-F | 5'-3'aaaGGTACCATGGTGAGCAAGGGCGA |
|  |  | GFP-ATG8e-Sac1-R | 5'-3'tttGAGCTCTTAGATTGAAGAAGCACCGAA |
| GFP-ATG8h-1776 |  | GFP-ATG8h-EcoR1-F | 5'-3'aaaGAATTCATGGTGAGCAAGGGCGAGG |
|  |  | GFP-ATG8h-R | 5'-3'GACAACAATCCCCATCCCGGGGAATTCG |
|  |  | ATG8h-GFP-F | 5'-3'CGAATTCCCCGGGATGGGGATTGTTGTC |
|  |  | ATG8h-GFP-Sac1-R | 5'-3'tttGAGCTCTTAGCCGAAAGTTTTCTCGGTG |
| GFP-ATG8i-1776 |  | GFP-ATG8i-EcoR1-F | 5'-3'aaaGAATTCATGGTGAGCAAGGGCGAGG |
|  |  | GFP-ATG8i-R | 5'-3'TCCTTGAACGATTTCATCCCGGGGAATTCG |
|  |  | ATG8i-GFP-F | 5'-3'CGAATTCCCCGGGATGAAATCGTTCAAGGA |
|  |  | ATG8i-GFP-Sac1-R | 5'-3'tttGAGCTCTCAACCAAAGGTTTTCTCACTGCTA |
| CLC2-MBP-1776 |  | CLC2-Kpn1-F | 5'-3'aaaGGTACCATGTCTGCCTTTGAAGACGAT |
|  |  | CLC2-MBP-R | 5'-3'CTTCTTCGATTTTCATCCCACCACCGGATC |
|  |  | MBP-CLC2-F | 5'-3'GATCCGGTGGTGGGATGAAAATCGAAGAAG |
|  |  | MBP-CLC2-Sac1-R | 5'-3tttGAGCTCTCAAGTCTGCGCGTCTTTC |
